# Rescue schizophrenia-related phenotypes caused by *Setd1a* deficiency by histone demethylase inhibitors

**DOI:** 10.64898/2026.08.03.742543

**Authors:** Yiqiong Liu, Guoguang Xie, Shan Jiang, Chengjie Zhou, Chao Zhang, Jun Qi, Ed Scolnick, Morgan Sheng, Yi Zhang

## Abstract

Schizophrenia (SCZ) is a genetically complex neuropsychiatric disorder in which rare loss-of-function mutations in the histone methyltransferase SETD1A confer substantial risk. Although SETD1A haploinsufficiency had been linked to morphological, synaptic and behavioral abnormalities in the prefrontal cortex, whether and how SETD1A coordinates transcriptional and functional programs across different brain regions remains unknown. Here, we delineate the brain region-specific effects of SETD1A-associated dysfunction using conditional *Setd1a* knockout mice. We find that the dorsal striatum (dStr) and mediodorsal thalamus (MD) exhibit distinct transcriptomic and neuronal alterations to those in the PFC, and show transcriptomic enrichment for other SCZ risk genes. Loss of *Setd1a* in the dStr or MD drives selective vulnerability in key behavioral assays, suggesting important roles for these brain regions in the etiology of SCZ. By screening 6 existing H3K4 demethylase inhibitors, we identify the LSD1 (KDM1A) inhibitor TAK-418 as a potent modulator capable of restoring H3K4me3 levels and gene expression, as well as rescuing synaptic and SCZ-like behavioral phenotypes in the *Setd1a*^+/−^ mice. Thus, our work provides a mechanistic link between high-penetrance SETD1A variants and region-specific brain dysfunction, establishing a framework for connecting rare loss-of-function variation in chromatin regulators to multidimensional neuropsychiatric phenotypes.

## Main

Schizophrenia (SCZ) is a severe and heterogeneous psychiatric disorder characterized by positive, negative, and cognitive symptoms, including delusions, hallucinations, anhedonia, and sensorimotor gating deficits^1,2^. Currently, the most widely used treatment for SCZ is antipsychotic-based medications^3,4^, which only provide limited relief for the negative and cognitive deficits that persist as lifetime features of the illness^5^. Consequently, more effective and mechanistic-based therapies are urgently needed^6^.

Accumulating evidence indicates that SCZ is a heterogeneous disease with diverse symptoms. The symptoms arise from dysfunction across multiple interconnected brain regions rather than isolated cortical abnormalities^7,8^. Thus, elucidating the brain region-specific mechanisms underlying key features of SCZ, such as anhedonia and sensorimotor-gating deficits, is essential for identifying molecular targets for therapeutic development.

Recent large-scale genetic studies, including the Schizophrenia Exome Sequencing Meta-Analysis (SCHEMA) consortium, have identified SETD1A as one of the most penetrant risk genes for SCZ, with loss-of-function (LoF) mutations conferring a substantial risk and providing critical mechanistic insights into SCZ pathogenesis^9^. Haploinsufficiency of SETD1A has been shown to impair gene transcription, neuronal morphology, synaptic transmission, and cognitive function in both animal and human stem cell models^10–14^. However, most prior studies have focused on the prefrontal cortex^15–17^, leaving unresolved question of how SETD1A deficiency impacts other brain regions and contributes to the diverse behavioral phenotypes.

As a histone methyltransferase, *SETD1A* encodes a core component of the COMPASS complex that catalyzes mono-, di-, and trimethylation of histone H3 lysine 4 (H3K4me1/2/3) ^18^, modifications that mark transcriptionally active enhancers and promoters^19^. LoF mutations in SETD1A disrupt these methylation patterns, reshaping the epigenetic landscape and leading to widespread transcriptional dysregulation associated with increased SCZ risk. Yet, the broader transcriptomic effects of SETD1A deficiency, the specific brain regions most vulnerable to its loss, and the links between the gene expression alterations to behavioral dysfunction remain poorly understood.

Epigenetic modifications, including histone methylation, play a pivotal role in neuronal development, synaptic plasticity, and circuit function^20–22^. Dysregulation of epigenetic modifications has been increasingly implicated in the pathophysiology of major psychiatric disorders, including SCZ^23^, depression^24^, and bipolar disorder^25^. Unlike genetic mutations, epigenetic modifications are reversible^26,27^, offering a promising avenue for therapeutic intervention. In recent years, small molecule epigenetic modulators, such as histone deacetylase (HDAC) inhibitors (e.g., valproate, sodium butyrate)^28^ and LSD1 demethylase inhibitors (TAK-418, TAK-488, ORY-1001) have been shown to be capable of rescuing social and memory deficits in animal models^11,29^. These findings raise the possibility of correcting transcriptional and synaptic abnormalities underlying SETD1A deficiency-associated SCZ by modulating H3K4 methylation.

To address the brain region-specific vulnerability to SETD1A insufficiency and their contributions to behavioral phenotypes, we characterized both *Setd1a* heterozygous (*Setd1a*^+/-^) and conditional knockout (*Setd1a*^flox/flox^) mouse models. By integrating molecular, neuronal, and behavioral analyses, we mapped alterations across multiple SCZ-relevant brain regions and identified pronounced deficits in the striatum and thalamus. Building on the concept of reversible epigenetic dysregulation, we explored pharmacological inhibition of histone H3K4 demethylases to rescue the abnormalities. Through systematic screening, we identified the LSD1 inhibitor TAK-418 as a potent chemical that restores H3K4 methylation, normalizes transcription and synaptic function, as well as ameliorates SCZ-like behaviors of *Setd1a*^+/-^ mice. Thus, our study defines a regionally resolved epigenetic framework linking SETD1A deficiency to SCZ pathophysiology and establishes the histone demethylase inhibitor TAK-418 as a potential therapeutic agent for SCZ caused by SETD1A deficiency.

## Results

### Region-specific deletion of *Setd1a* reveals distinct behavioral deficits

Schizophrenia (SCZ) is characterized by widespread dysregulation across multiple brain regions, including reduced parvalbumin-positive neurons in the cerebral cortex and hippocampus^30^, reduced prefrontal cortex (PFC) neuronal activity^31^, abnormalities in the dopamine (DA) system in the striatum^32,33^, accompanied by structural and metabolic abnormalities^34^. Given that *Setd1a* encodes a histone methyltransferase essential for transcriptional regulation and neuronal development^35^, its haploinsufficiency is expected to exert region-specific effects on brain function. Although *Setd1a* is broadly expressed throughout the adult mouse brain, with highest levels in the neocortex, prior studies have focused almost exclusively on the PFC^11,12^. To delineate the broader functional landscape of *Setd1a* deficiency, we systematically examined its molecular, neuronal, and behavioral consequences across major SCZ-relevant brain regions.

To confirm the brain regions mentioned above are indeed involved in SCZ-related behaviors, we first performed pre-pulse inhibition test (PPI) assays, one of the most reliable SCZ behavior tests^36^, coupled with c-Fos staining (Extended Data Fig.1a). This analysis revealed robust neuronal activation in the medial prefrontal cortex (mPFC), dorsomedial striatum (dStr), medial nucleus accumbens (NAcMed), and mediodorsal thalamus (MD) (Extended Data Fig.1a,b), supporting their involvement in the behaviors affected in SCZ.

To facilitate the study of brain region-specific function of *Setd1a*, we generated a conditional *Setd1a* knockout (*Setd1a*^flox/flox^) mouse line with *loxP* sites flanking exons 14-16, which encodes the SET domain of *Setd1a* (Extended Data Fig.1c,d). To achieve brain region-restricted deletion, adeno-associated viruses (AAVs) encoding Cre-mCherry or mCherry (control) were stereotaxically injected into SCZ-relevant regions (Fig.1a), including mPFC, dStr, NAc, MD, hippocampus (HPC), and cerebellar cortex (CB). Single-molecule fluorescence in situ hybridization (smFISH) revealed no detectable *Setd1a* mRNA expression in nearly all Cre-mCherry-expressing cells (∼99%), confirming efficient *Setd1a* depletion in virus injected area (Fig.1b). The expression of Cre-mCherry in the virus injected sites was confirmed by *post hoc* histological examination (Fig.1c).

**Fig. 1:**
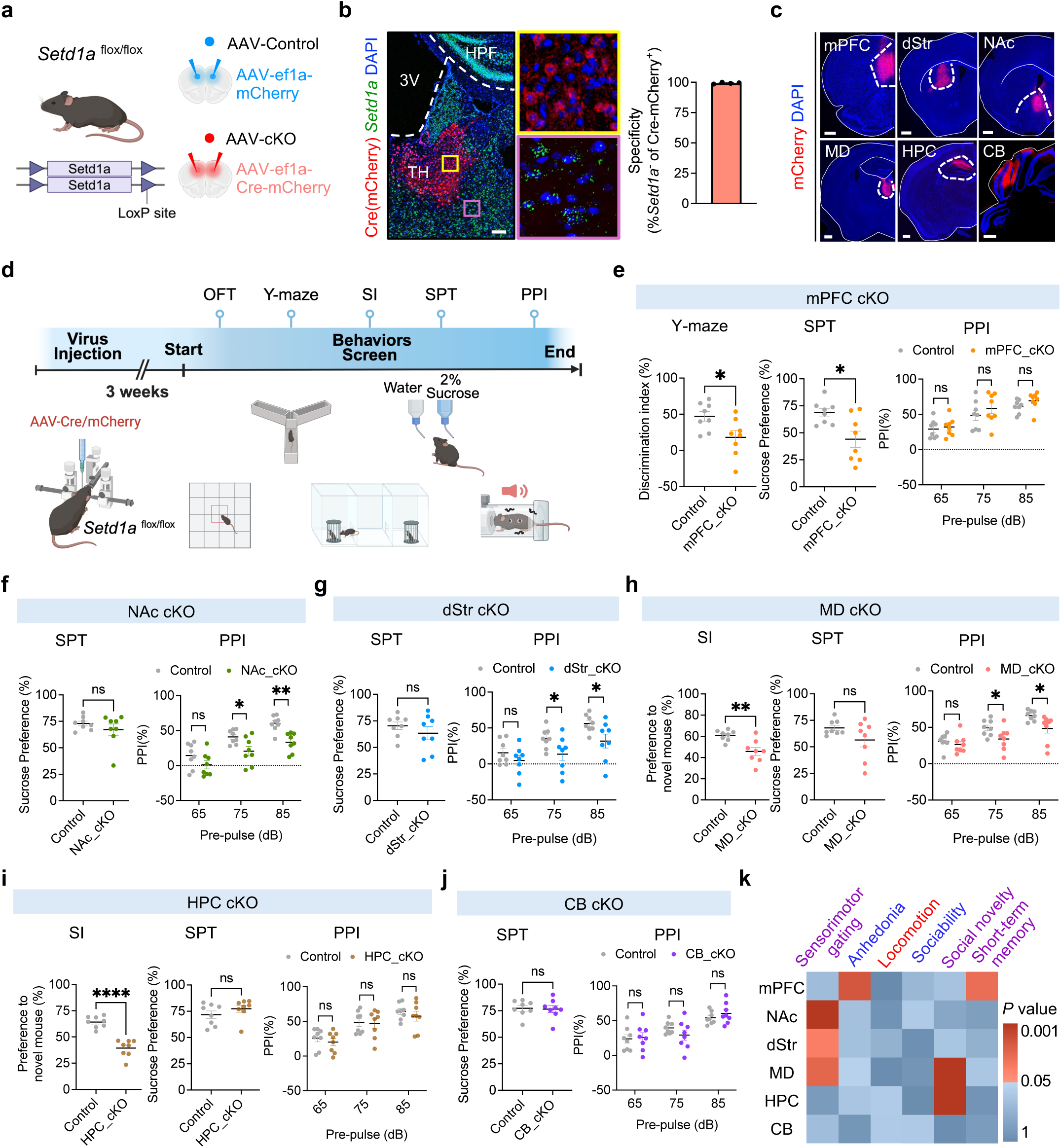
Brain region-specific deletion of *Setd1a* causes distinct behavioral deficits. **a**, Schematic of *Setd1a*^flox/flox^ mice generation, with viral injection sites in the medial prefrontal cortex (mPFC) shown as an example. **b**, Left, smFISH confirms that *Setd1a* is efficiently knocked out following Cre-expressing AAV injection. Middle, enlarged views of yellow and purple boxed regions. Right, quantification of the percentage of *Setd1a*^-^ cells among Cre-mCherry positive cells (n=4 sections from3 mice). Scale bar, 200 μm. 3V, third ventricle. TH, thalamus. HPF, hippocampal formation. **c**, Representative images of Cre-mCherry expression in the viral injected sites of medial prefrontal cortex (mPFC), dorsal striatum (dStr), nucleus accumbens (NAc), medial dorsal thalamus (MD), hippocampus (HPC), or cerebellar cortex (CB), respectively. Scale bar, 500 μm. **d**, Experimental timeline for behavioral screening of *Setd1a* conditional knockout (cKO) mice. **e**, Left, discrimination index in the Y maze test of mPFC control and *Setd1a* cKO mice. Middle, percentage of sucrose consumption of total liquid intake. Right, pre-pulse inhibition (PPI) values at three pre-pulse intensities (65, 75 and 85 dB) (n=8 per group). **f**, Left, percentage of sucrose consumption of NAc control and *Setd1a* cKO mice. Right, PPI values at three different pre-pulse intensities (n=8 per group). **g**, Left, percentage of sucrose consumption of dStr control and *Setd1a* cKO mice. Right, PPI values at three different pre-pulse intensities (n=8 per group). **h**, Left, social preference for the chamber containing a novel mouse of MD control and *Setd1a* cKO mice in the three-chamber social interaction test (SI). Middle, percentage of sucrose consumption of total liquid intake. Right, PPI values at three different pre-pulse intensities (n=8 per group). **i**, Left, social preference for the chamber containing a novel mouse of HPC control and *Setd1a* cKO mice in SI. Middle, percentage of sucrose consumption of total liquid intake. Right, PPI values at three different pre-pulse intensities (n=8 per group). **j**, Left, percentage of sucrose consumption of CB control and *Setd1a* cKO mice. Right, PPI values at three different pre-pulse intensities. (n=8 per group) **k**, Heatmap summary of the behavioral performance of the *Setd1a* cKO mice. The functional domains are color-coded: purple, cognitive; blue, negative-like; red, positive-like behaviors. Data were presented as means ± SEM. Each dot represents the result from an individual mouse. Two-tailed unpaired Student’s *t* test. ns, not significant; \**P* < 0.05, \*\**P* < 0.01, \*\*\**P* < 0.001, \*\*\*\**P* < 0.0001.

We next assessed the behavioral consequences of region-specific *Setd1a* deletion by subject the mice to a battery of SCZ-related behavioral assays, including the open field test (OFT), Y-maze test, three chamber social interaction test (SI), sucrose preference test (SPT), and PPI test (Fig.1d). mPFC specific conditional knockout (mPFC-cKO) mice exhibited reduced novel arm exploration in the Y-maze and decreased sucrose preference (Fig.1e), resembling cognitive and motivational impairments characteristic of SCZ’s negative and cognitive symptoms. No significant changes were observed in locomotion, social preference, or PPI (Fig.1e and Extended Data Fig.2a). In contrast, *Setd1a* deletion in the NAc (NAc-cKO) and dStr (dStr-cKO) selectively impaired PPI (Fig.1f,g), but not other behaviors (Fig.1f, g and Extended Data Fig.2b,c), reflecting impaired sensorimotor gating. Deletion in the MD (MD-cKO) disrupted both social novelty preference and PPI (Fig.1h), but not other behaviors (Fig.1h and Extended Data Fig.2d), suggesting a role in integrating behavioral responses relevant to social and attentional regulation. HPC specific knockout (HPC-cKO) altered social novelty preference only (Fig.1i and Extended Data Fig.2e), while cerebellar deletion (CB-cKO) had no detectable behavioral effect (Fig.1j and Extended Data Fig.2f).

Together, these findings demonstrate that Setd1a is required in a brain region-specific manner to maintain specific behavioral domains relevant to SCZ, with executive and reward processing in the mPFC, sensorimotor gating in the striatum and thalamus, and social affective regulation in the hippocampus and thalamus (Fig.1k). The regional selectivity of these phenotypes suggests that SETD1A haploinsufficiency contributes to the heterogeneous symptoms of SCZ through brain region-specific mechanisms, rather than a uniform, brain-wide effect.

### *Setd1a* loss disrupts neuronal activity dynamics underlying sensorimotor gating and reward processing

The region-specific behavioral alterations observed in *Setd1a* conditional knockout mice suggest that *Setd1a* contributes to multiple functional domains relevant to SCZ. Given the prominent involvement of the cortico-striatal-thalamic (CST) network in sensorimotor gating and reward processing, two key cognitive and negative symptoms in SCZ^37^, we next examined the effects of *Setd1a* deficiency on the neuronal activity of these regions.

To this end, we performed multi-site fiber photometry during PPI test (Fig.2a). Neuronal calcium activity was visualized using AAV vectors carrying a Cre-dependent FLEX cassette to drive expression of the genetically encoded calcium indicator GCaMP7s. To ensure comparable infection efficiency and allow precise genotype-dependent comparisons, AAV mixtures encoding Cre and GCaMP7s was co-injected into matched subregions of the mPFC, NAc, dStr, and MD in wide-type (WT) and *Setd1a^+/-^* mice (Fig.2c).

**Fig. 2:**
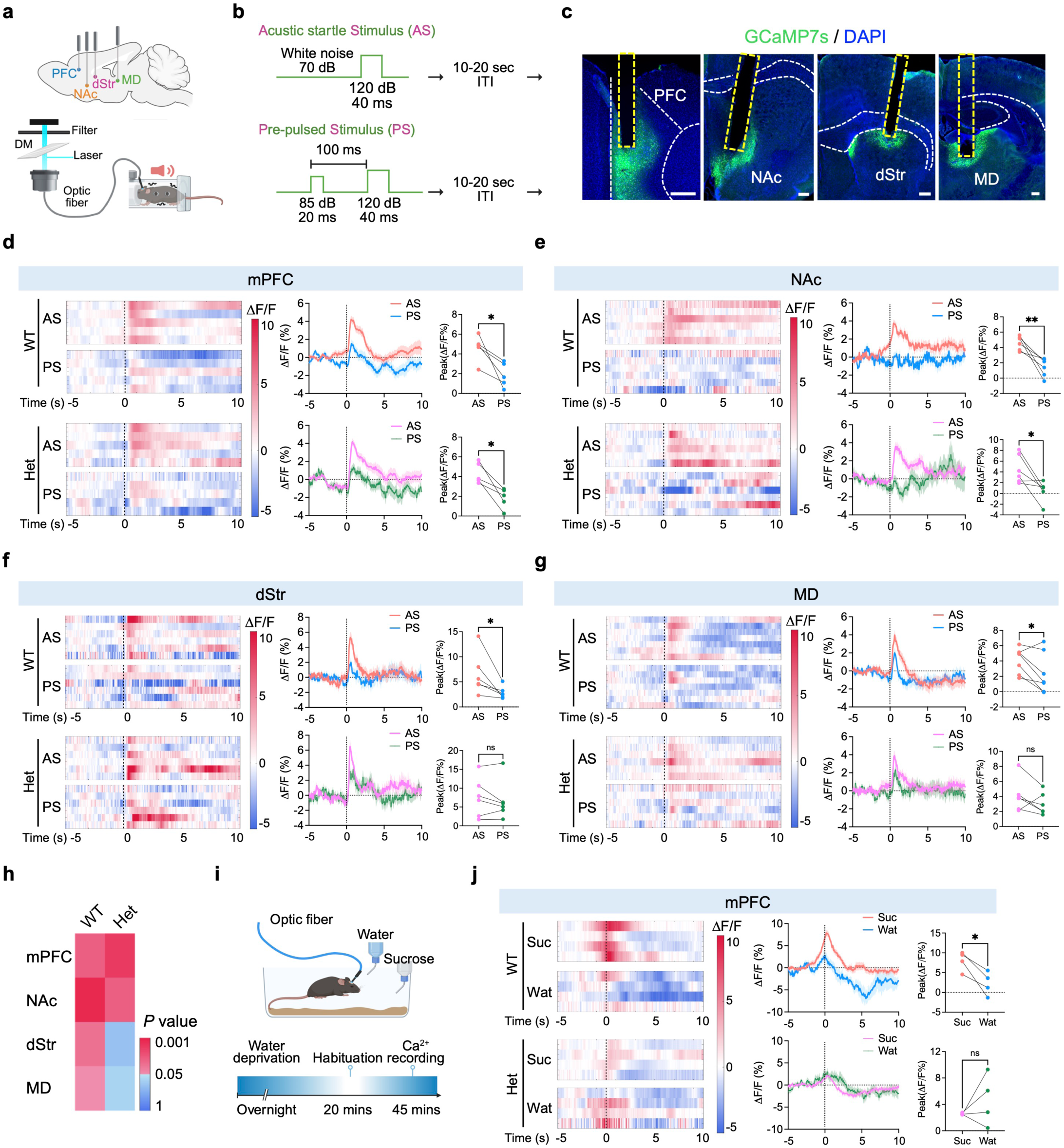
Setd1a loss disrupts neuronal activity dynamics underlying sensorimotor gating and reward processing. **a**, Schematic diagram illustrating multi-site fiber photometry setup for recording calcium activities during PPI. **b**, Illustration of the two types of stimuli. **c**, Representative images showing viral expression and optic fiber placement in four brain regions (mPFC, NAc, dStr and MD). Green, GCaMP7s; blue, DAPI. Yellow dotted line, fiber track. Scale bar, 200 µm. **d**, Left, heat maps of averaged ΔF/F Ca^2^ responses in the mPFC of individual WT (top) or *Setd1a^+/-^* mice (bottom) aligned to stimulus onset (each row represents one mouse). Middle, averaged peri-stimulus traces of Ca^2^ signals of AS and PS. Dashed line, event onset. Right, quantification of averaged peak ΔF/F values (dot plot). Two-tailed, paired *t*-test. n=5 mice. **e-g,** Same as in **d**, but for NAc (**e**), dStr (**f**) and MD (**g**). Two-tailed, paired *t*-test. n=6-7 mice. **h**, Summary of the difference between AS and PS in WT or *Setd1a^+/-^* mice across the four different brain regions. *P* values are color-coded. **i**, Schematic of the recording setup in freely behaving mice during the sucrose preference test. **j**, Left, heat maps of averaged ΔF/F Ca^2^ responses in the mPFC of individual WT (top) or *Setd1a^+/-^* mice (bottom) aligned to the onset of drinking (each row represents one mouse). Middle, averaged peri-stimulus traces of Ca^2^ signals. Dashed line, event onset. Right, quantification of averaged peak ΔF/F values (dot plot). Two-tailed, paired *t*-test. n=4 mice. Data are means ± SEM. Each dot represents one mouse. ns, not significant; \**P* < 0.05, \*\**P* < 0.01.

Consistent with c-Fos mapping (Extended Data Fig.1a,b), acoustic startle stimuli (AS) evoked robust calcium transients in all four regions of both genotypes (Fig.2d-g, left). To directly compare neuronal responses to the acoustic startle tone (AS; 120 dB) and pre-pulsed stimulus (PS; 85 dB pre-pulse + 120 dB startle tone) (Fig.2b), calcium traces were aligned to the onset of the tone. We observed that WT mice showed significant higher peak amplitude of AS response than PS response across all these brain regions, consistent with pre-pulse induced motor inhibition and the CST network engagement during sensorimotor gating (Fig.2d-g, top). In the mPFC and NAc, *Setd1a^+/-^*mice also displayed lower peak amplitude of PS compared to AS, similar to that of the WT mice (Fig.2d, e, bottom), indicating that Setd1a haploinsufficiency in mPFC and NAc do not impair neuronal dynamics during PPI. The result of mPFC align with behavioral screening data (Fig.1e), which showed preserved PPI performance. While NAc *Setd1a* cKO showed PPI deficit (Fig.1f), it suggests a potential gene dose-dependent contribution of *Setd1a* in NAc-mediated sensorimotor gating.

In contrast, in the dStr and MD, WT mice exhibited strong calcium transients following the AS and blunted this response upon PS, reflecting normal pre-pulse modulation (Fig.2f, g, top), whereas *Setd1a*^+/−^ mice diminished the differential responses of AS and PS (Fig.2f, g, bottom), suggesting Setd1a haploinsufficiency disrupted signal integration in motor gating. Together, these results demonstrate that while mPFC and NAc activity remains largely intact, *Setd1a* loss selectively dampens the pre-pulsed stimulus-evoked calcium dynamics in the dStr and MD that collectively mediate the sensorimotor gating deficits observed in *Setd1a^+/-^*mice^16^ (Fig.2h).

To further explore Setd1a-dependent neural activity that correlates to reward processing, we recorded neuronal calcium dynamics in the mPFC of WT and *Setd1a^+/-^* mice during sucrose preference test since only mPFC-cKO mice exhibited a deficit (Fig.1e). Following overnight water deprivation, mice were allowed free access to water or sucrose solutions in their home cages (Fig.2i). In both genotypes, neuronal activity increased as animals approached the water bottle, peaked during consumption, and declined thereafter, reflecting motivation and satiety-associated dynamics upon drinking behaviors (Fig.2j, left). To assess neural activity that correlates to sucrose preference, we compared calcium signals during sucrose versus water consumption. In WT mice, mPFC neurons showed markedly enhanced activity during sucrose intake, with significantly higher peak amplitudes relative to that of the water (Fig.2j, top). In contrast, *Setd1a*^+/−^ mice exhibited blunted and comparable responses to sucrose and water (Fig.2j, bottom), consistent with their reduced sucrose preference (Fig.1e, middle).

Together, these results show that Setd1a haploinsufficiency alters neuronal activity in a region-specific manner. Reduced responsiveness in the dorsal striatum and thalamus corresponds to sensorimotor gating deficits, while diminished mPFC activity during reward consumption correlates with motivational impairments. Our findings link alterations of neuronal activity dynamics to SCZ-related behavioral changes in response to Setd1a loss.

### Setd1a haploinsufficiency induces brain region-specific transcriptomic alterations

Given the brain region-selective behavioral and neuronal activity phenotypes, we next sought to define the molecular alterations underlying Setd1a deficiency in the SCZ-relevant regions. To this end, we performed RNA sequencing (RNA-seq) analysis on mPFC, NAc, dStr and MD (Fig.3a) from *Setd1a^+/−^* and their WT littermates at 8 weeks of age, corresponding to the regions exhibiting behavioral and neuronal activity phenotypes. Replicates exhibit high correlations, confirming the data quality (Extended Data Fig.3a-d). Comparative analysis identified different numbers of differentially expressed genes (DEGs) across all four regions (Fig.3b; Supplementary Table 1) with NAc showed the largest transcriptomic shift (691 up and 443 down), while mPFC displayed the least changes (147 up and 150 down). Cross-region comparison revealed that most DEGs were region-specific, with only 280 DEGs were shared between any two regions and only 9 were shared among three regions (Fig.3c). Interestingly, comparison of the DEGs with those of the SCZ risk genes identified by GWAS and SCHEMA revealed relative enrichment in NAc, dStr and MD (Fig.3d). These results indicate that Setd1a loss exhibits brain region-specific transcriptomic effects.

**Fig. 3:**
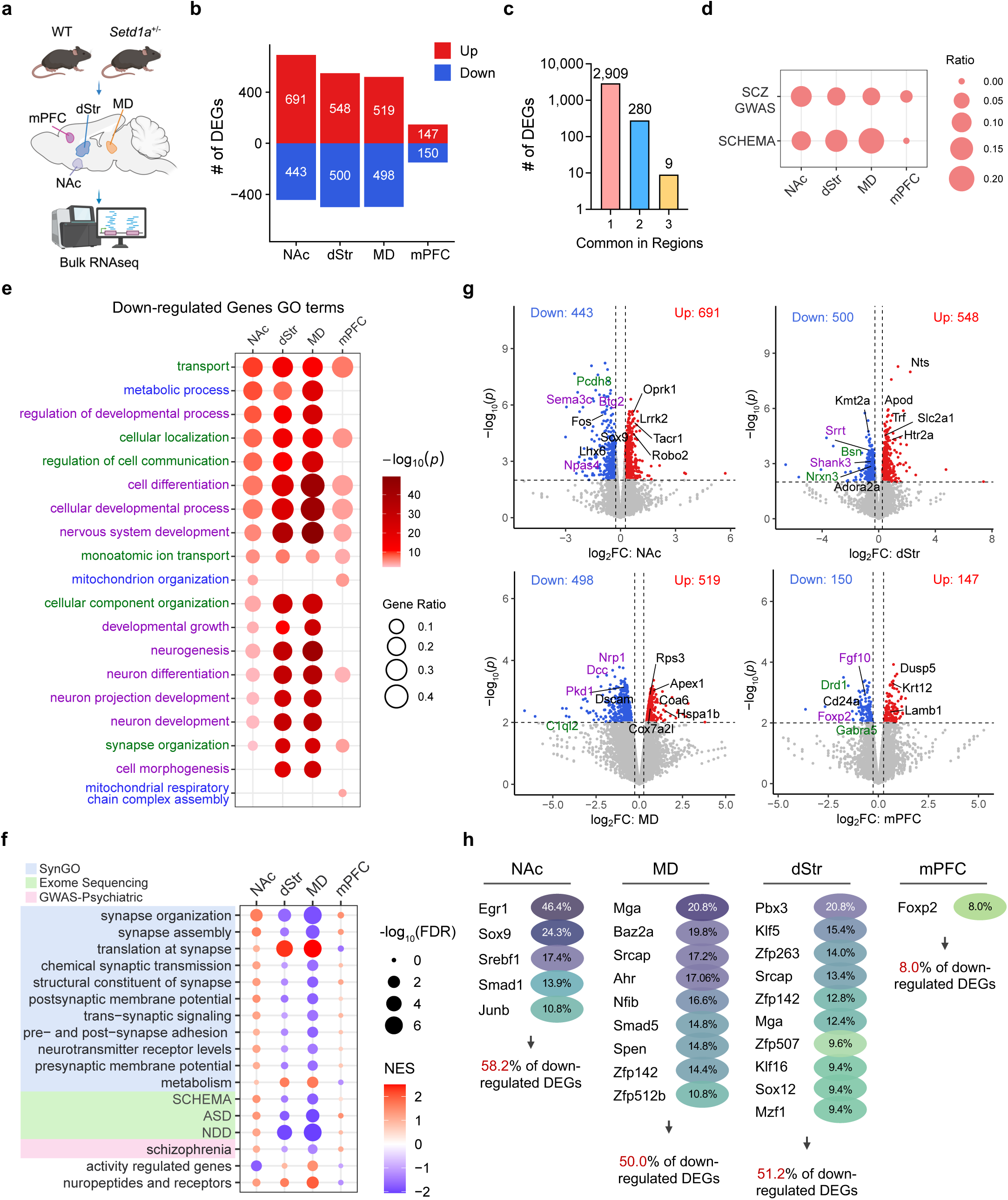
Setd1a haploinsufficiency induces brain region-specific transcriptome alterations. **a**, Schematic diagram of bulk RNA-seq of multiple brain regions in WT and *Setd1a^+/-^* mutant mice. n=2 mice per genotype. **b**, Number of DEGs in the indicated brain regions. Blue and red bars represent down- and up-regulated DEGs, respectively. Cutoff: *p* <0.01 with |fold change| >1.2 **c**, Number of DEGs showing brain region-specific or common in more than one brain region. **d**, Dot plot showing the ratio of differentially expressed risk genes associated with SCZ-GWAS and SCHEMA. **e**, Dot plot showing the Gene Ontology (GO) terms enriched for the down-regulated DEGs of four different brain regions. The −log_10_(*p*) of GO terms are color-coded. Purple, nervous system development processes; Green, cellular organization; Blue, cellular energy metabolism. **f**, GSEA analysis of DEGs with SynGo reveals enrichment of psychiatric diseases, activity-regulated gene set and neuropeptides. **g**, Volcano plots of transcriptomic changes found in the bulk RNA-seq analysis of the indicated brain regions of *Setd1a^+/-^* mice. Blue and red dots represent down- and up-regulated DEGs, respectively. Genes in purple or green color are correlated with terms. Log_2_FC, log_2_Fold Change. **h**, Transcription factor (TF) enrichment analysis showing target genes of the denoted TFs and their percentage overlap between down-regulated DEGs in NAc, MD, dStr and PFC.

To gain insight into the biological processes affected by Setd1a haploinsufficiency, we performed Gene Ontology (GO) enrichment analysis. We found that down-regulated genes were significantly enriched in pathways related to nervous system development (highlight in purple), cellular organization (highlight in green), and cellular energy metabolism (highlight in blue) (Fig.3e). These processes are consistent with findings from transcriptomic and proteomic studies using postmortem SCZ patient brains ^38,39^. Gene set enrichment analysis (GSEA) using the refined SynGO terms^40^ further revealed disruptions in both pre- and postsynaptic processes. Of note, DEGs of dStr and MD are negatively associated with synapse organization, membrane potential, synapse adhesion (Fig.3f). Moreover, DEGs also associate with autism disorder (ASD) and neurodevelopmental disorder (NDD). In contrast, upregulated genes were enriched for GO terms such as “cellular component organization”, “system development” and “cellular respiration” (Extended Data Fig.3e), suggesting compensatory or maladaptive remodeling responses.

Among the downregulated genes, several were associated with nervous system development (highlighted in purple), such as *Sema3c*, *Btg2*, *Npas4* in NAc; *Srrt*, *Shank3* in dStr; *Nrp1*, *Pkd1, Dcc* in MD; *Fgf10, Foxp2* in mPFC. Some were implicated in synaptic organization (highlighted in green), including *Pcdh8* in NAc; *Bsn*, *Nrxn3* in dStr; *C1ql2* in MD; *Drd1*, *Gabra5* in mPFC (Fig.3g). Many of these DEGs exhibited regional selective effect. For instance, *Foxp2*, a transcription factor enriched in deep-layer PFC neurons, was selectively downregulated in the mPFC of *Setd1a^+/−^* mice, consistent with our previous single-cell RNA-seq result^16^ (Extended Data Fig.3f).

Setd1a loss additionally altered the expression of several key transcriptional regulators, including *Kmt2a* in the dStr, implying that some of the transcriptional alterations might be caused by the changes of these transcriptional regulators. To identify potential transcription factors (TFs) mediating these transcriptional alterations, we performed TF enrichment analysis based on orthogonal omics integration^41^. Among the TFs derived from the enrichment analysis, we focused on TFs that themselves are DEGs of *Setd1a* mutants and then identified the overlapped genes between the denoted TFs’ targeted genes and *Setd1a* mutants DEGs. The percentage (overlapping genes/total DEGs) represents the transcriptional alterations potentially mediated by each TF. We found that these TFs accounted for over 50% of the down-regulated DEGs (Fig.3h), and over 20% of the up-regulated DEGs in the NAc, MD, and dStr (Extended Data Fig.3g).

Among the down-regulated TFs, several are particularly relevant. For example, *Egr1* positively regulates genes involved in synaptic plasticity and learning^42^; *Ahr* promotes dendritic arborization and GABAergic neurons differentiation^43^; while *Pbx3* regulates neuronal differentiation, synaptic development, and neuronal excitability^44,45^. On the other hand, upregulated TFs included *Creb1,* a cAMP- and calcium-responsive activator that recruits CBP/p300 to activate gene expression^46^; *Smad5*, an intracellular signal transducer protein that is up-regulated in postmortem SCZ brain^47^; and *Cebpd*, a transcriptional activator involved in inflammation, metabolism, and stress response. The functional categories enriched among their targets, such as “response to stress” and “catabolic process”, are consistent with the broader transcriptional alterations of Setd1a loss (Extended Data Fig.3e,g).

Together, these results demonstrate that Setd1a orchestrates region-specific transcriptional programs that regulate neuronal development, synaptic function, and metabolic homeostasis across cortical and subcortical regions. Its deficiency triggers both direct and indirect transcriptional dysregulation, explaining how alteration of an epigenetic factor can give rise to diverse neurobiological phenotypes characteristic of SCZ.

### Pharmacological reversal of Setd1a haploinsufficiency via histone demethylase inhibition

Since SETD1A encodes a lysine methyltransferase that could catalyze mono-, di-, and tri-methylation on H3K4 to regulate transcription, pharmacological inhibition of H3K4 demethylases, such as the LSD1 and KDM5 family members, may compensate the loss of H3K4 methylation due to *Setd1a* heterozygosity to correct the transcriptional dysregulation. Consistent with this notion, a previous study showed that the LSD1 inhibitor ORY-1001 could rescue cognitive and morphological deficits of *Setd1a*^+/-^ mice^11^. To identify more effective H3K4 demethylase inhibitors capable of rescuing negative and cognitive deficits (anhedonia and impaired sensorimotor gating) compensate for *Setd1a* heterozygosity, we screened several KDM5 inhibitors (KDM5-C70, Cpd-48, CPI-455, JQKD82) and LSD1 inhibitors (TAK-418, ORY-1001) with reported brain penetrance and favorable pharmacological profiles^48,49^.

We first tested the ability of these inhibitors to rescue the morphological defects of cultured primary cortical neurons derived from *Setd1a*^+/−^ mice. To this end, cultured neurons were treated from days in vitro (DIV) 3-10 with different concentrations of each compound, and neurite outgrowth was quantified every two days (Fig.4a). Among the inhibitors tested, JQKD82 (0.1-1 µM) and TAK-418 (0.3-3 µM) could restore the *Setd1a*^+/−^ neuron branch numbers to that of the WT levels (Fig.4a,b), whereas other compounds failed to rescue or even caused neuron death (Extended Data Fig.4). These results suggest that both JQKD82 and TAK-418 could compensate for the loss of neurite outgrowth in the *Setd1a*^+/−^ cortical neurons.

**Fig. 4:**
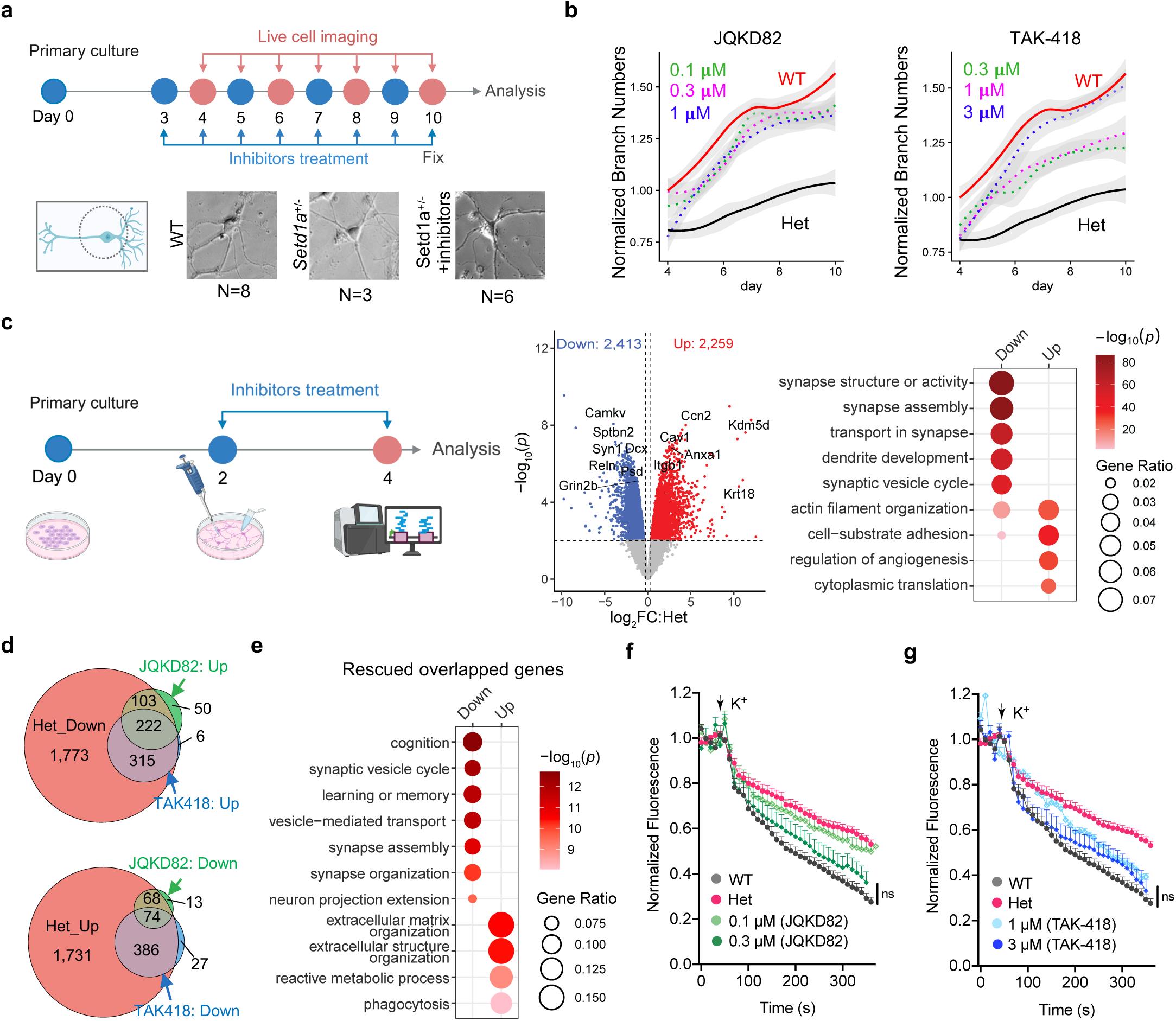
Pharmacological reversal of Setd1a haploinsufficiency via histone demethylase inhibition *in vitro*. **a**, Top, schematic of the experimental timeline for morphological analysis. Bottom, representative images of cultured neurons at DIV10. N indicates the primary branch numbers. **b**, Quantification of normalized neurite numbers compared to WT group across 10 days *in vitro*. Dotted lines indicate inhibitor treatment groups; colors denote experimental conditions. n= 50 cells per group. **c**, Left, experimental design for RNA-seq after inhibitor treatment. Middle, Volcano plot showing transcriptomic changes comparing cultured WT and *Setd1a^+/-^* neurons. Right, dot plot showing the GO terms enriched for the down-regulated DEGs in *Setd1a^+/-^* neurons. **d**, Top, overlap of down-regulated genes in *Setd1a^+/-^* neurons with upregulated genes after JQKD82 or TAK-418 treatment; Bottom, overlap of up-regulated genes in *Setd1a^+/-^* neurons with down-regulated genes after JQKD82 or TAK-418 treatment. **e**, GO enrichment analysis of overlapping DEGs of JQKD82 and TAK-418 treatments. GO terms are color-coded by –log (*p*) value. **f**, Time-lapse recording of K^+^-induced exocytosis in cortical (left) and striatal (right) neurons. The FM1-43 signals at different time points were plotted. The arrow indicates the time when 50 mM K^+^ was applied. WT, neurons from WT mice; *Setd1a*^+/−^, neurons from *Setd1a*^+/−^ mice; JQKD82_0.1 μM and JQKD82_0.3 μM, neurons from *Setd1a*^+/−^ mice treated with 0.1 μM or 0.3 μM JQKD82, respectively. The arrow indicates the time when 50 mM K^+^ was applied. One-way ANOVA test with Holm-Šídák’s multiple comparisons test. n=30 neurons per group. **g**, Same as **e**but for TAK-418 treatment. One-way ANOVA test with Holm-Šídák’s multiple comparisons test. n=30 neurons per group.

We next asked whether JQKD82 or TAK-418 could rescue the transcriptional defects of *Setd1a*^+/−^ neurons. To this end, we performed RNA-seq on cultured primary cortical neurons treated with JQKD82 (1 μM), TAK-418 (3 μM), or ORY-1001(1 nM) from DIV2-DIV4 (Fig.4c and Extended Data Fig.5a). We included ORY-1001 because a previous study reported its phenotypical rescue effect^11^. Comparative analysis revealed that *Setd1a*^+/−^ cortical neurons harbor 2,413 down-regulated genes that are enriched in synaptic functions, while the 2,259 up-regulated genes are linked to actin organization and angiogenesis (Fig.4c; Supplementary Table 2). Interestingly, TAK-418 treatment rescued 537 down-regulated and 460 up-regulated genes in *Setd1a*^+/−^ neurons; JQKD82 rescued 325 down-regulated and 142 up-regulated genes, respectively (Fig.4d). Although ORY-1001 induced widespread transcriptional changes (4,363 DEGs), its overlap with *Setd1a*^+/−^ DEGs was minimal (Extended Data Fig.5b,c). Since ORY-1001 failed to rescue the neurite outgrowth or transcriptional alterations of *Setd1a*^+/−^ cultured cortical neurons, this inhibitor was not analyzed further. Since TAK-418 and JQKD82 rescued 640 downregulated genes with 222 in common (Fig.4d, top panel), we performed GO analysis of DEGs which revealed enrichment of cognitive and synaptic functions (Fig.4e; Extended Data Fig.5d,e). Similar analysis of the 74 commonly rescued up-regulated genes revealed the enrichment of extracellular matrix organization and metabolic process (Fig. 4d,e; Extended Data Fig.5d,e). These results indicate that both JQKD82 and TAK-418 could rescue the transcriptional defects of cultured *Setd1a*^+/−^ cortical neurons.

To determine whether transcriptional rescue leads to functional recovery, we measured exocytosis at DIV18 following inhibitor treatment initiated at DIV1. Both 0.3 μM JQKD82 and 3 μM TAK-418 treatment resulted in significant recovery of exocytosis in cultured *Setd1a*^+/−^ cortical neurons (Fig.4f, g), confirming rescue of synaptic function *in vitro*.

Collectively, the above results demonstrate that pharmacological inhibition of H3K4 demethylases, particularly TAK-418 and JQKD82, can effectively reverse the neuronal and synaptic abnormalities due to Setd1a haploinsufficiency, which highlights the potential of epigenetic modulation of H3K4 methylation as a therapeutic avenue for SETD1A-related SCZ.

### TAK-418 reverses behavioral abnormalities and restores H3K4me3-dependent transcriptional and synaptic function in *Setd1a*^+/−^ mice

Next, we evaluated whether these compounds can ameliorate SCZ-related behaviors *in vivo*. We found that three weeks of intraperitoneal (i.p.) administration of JQKD82 (70 mg/Kg, twice daily) to *Setd1a*^+/−^ mice restored their sucrose preference, but not the PPI deficits (Extended Data Fig.6). Extending the treatment to five weeks similarly failed to rescue PPI impairment (Fig.5a). Importantly, two weeks of oral TAK-418 treatment (1 mg/kg) rescued both sucrose preference and PPI defects of the *Setd1a*^+/−^ mice (Fig.5b). Neither treatment affected body weight, body temperature, nor locomotor activity (Fig.5c-e), supporting the potential of these inhibitors.

**Fig. 5:**
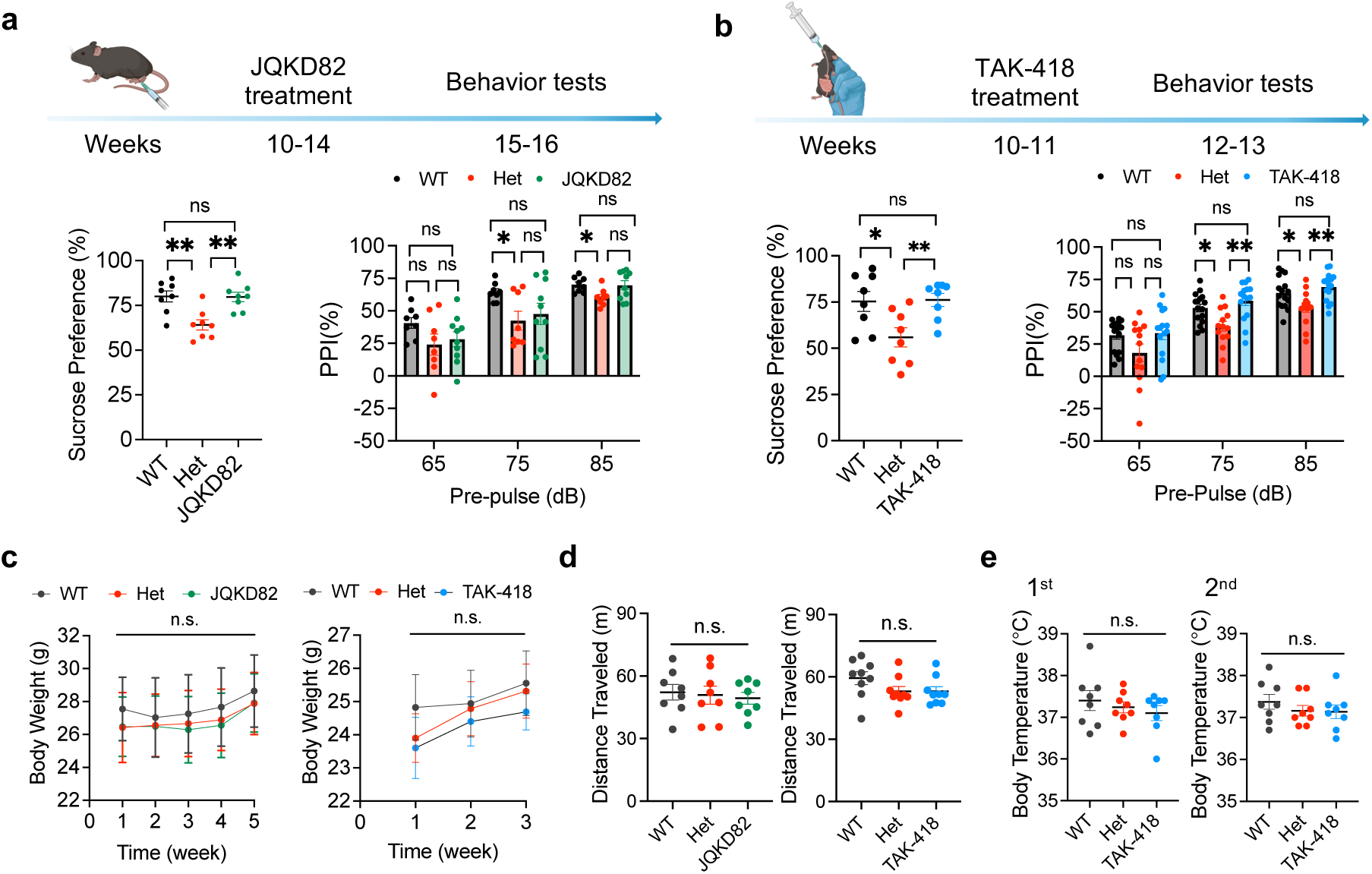
Histone demethylase inhibition partially rescues behavioral deficits in *Setd1a*^+/−^ mice. **a**, Behavioral assays following 5 weeks of JQKD82 treatment. Top, experimental timeline. Bottom, sucrose preference (left) and PPI (right) tests of WT, *Setd1a*^+/−^ and JQKD82 treated *Setd1a*^+/−^ mice. One-way ANOVA test with Tukey’s multiple comparisons test. n=8 per each group for sucrose preference test; n=8 for WT and Het groups, n=10 for JQKD82 group for PPI test. **b**, Behavioral tests after TAK-418 treatment. Top, experimental timeline. Bottom, sucrose preference (left) and PPI (right) tests of WT, *Setd1a*^+/−^ and TAK-418 treated *Setd1a*^+/−^ mice. One-way ANOVA test with Tukey’s multiple comparisons test. n=8 per each group for sucrose preference test; n=16 for WT, n=13 for Het groups, n=15 for TAK group for PPI test. **c**, No obvious effect of JQKD82 (left) or TAK-418 (right) administration on body weight. Two-way ANOVA. n=8 per group. **d**, Open field tests of total distance traveled after JQKD82 (left) or TAK-418 (right) treatment. One way ANOVA. n=8 per group (left); n=9 per group (right). **e**, Body temperature measurements after TAK-418 treatment for 1 week (left) and 2 weeks (right). One way ANOVA. Each dot represents the result from an individual mouse. n=8 per group. Each dot represents the result from an individual mouse. Data were presented as means ± SEM. n.s., not significant; \**P* < 0.05, \*\**P* < 0.01. Data were presented as means ± SEM. ns, not significant; \**P* < 0.05, \*\**P* < 0.01, \*\*\**P* < 0.001.

Given TAK-418 exhibits a stronger rescue effect on both transcriptome of cultured *Setd1a*^+/−^ cortical neurons and behavioral deficits of *Setd1a*^+/−^ mice compared to JQKD82 (Fig.4d, 5a,b), we next attempted to uncover the molecular mechanism by which TAK-418 might alleviate the behavioral deficits. To this end, we performed RNA-seq on isolated dStr, MD, NAc, and mPFC tissues from TAK-418 treated *Setd1a*^+/−^ mice (Extended Data Fig.7a). We then performed integrated transcriptome analyses on WT, *Setd1a*^+/−^, and *Setd1a*^+/−^ with TAK-418 treatment. We found that 369 out of the 443 (83.3%) down-regulated genes in the *Setd1a*^+/−^ NAc were rescued or partially rescued by the TAK-418 treatment (Fig.6a, b, left; Supplementary Table 3). Similar rescue or partial rescue effects were also observed in MD (67.9%), dStr (95.4%), and mPFC (94.7%) with many of the down-regulated genes recovered to levels of the WT (Fig.6a, b). For the up-regulated genes, TAK-418 treatment rescued most gene expression in the NAc, dStr, and mPFC, but only partially rescued in the MD (Fig.6a, b). GO enrichment analysis revealed that the rescued down-regulated genes are commonly enriched for cellular localization, and nervous system development, while those in MD and dStr were additionally enriched for synaptic organization and cell morphogenesis (Fig.6c). Similar analysis revealed that the rescued up-regulated genes were enriched for cellular component organization, system development and response to stress (Extended Data Fig.7b). Collectively, these data indicate that TAK-418 treatment results in restoration of the transcriptional programs disrupted by Setd1a deficiency, consistent with the previous GO analysis comparing WT and *Setd1a*^+/−^ mice (Fig.3e).

**Fig. 6:**
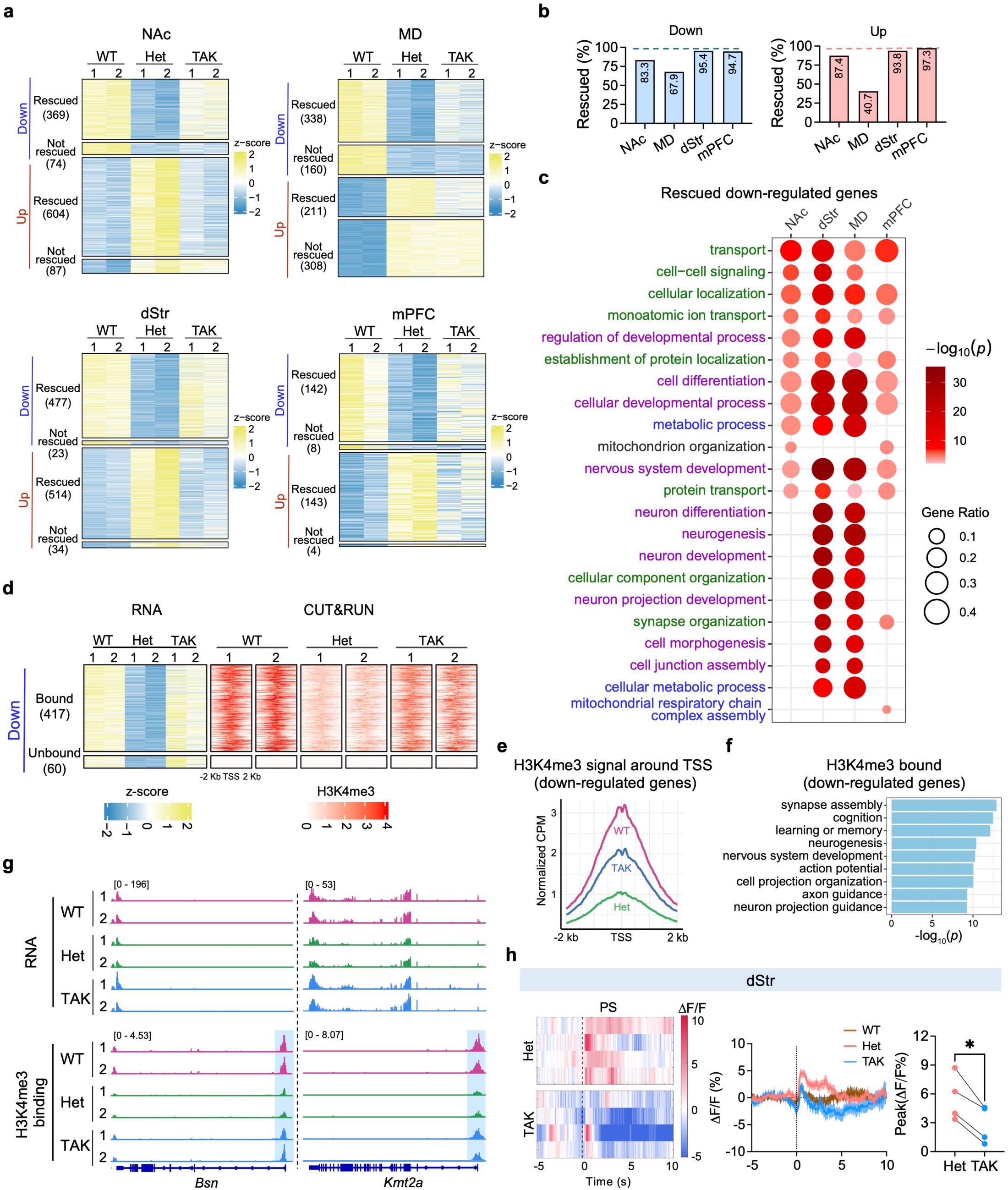
TAK-418 treatment restores gene expression and H3K4me3 occupancy in *Setd1a*^+/−^ mouse brain. **a**, Heatmaps showing the restoration of the z-scored expression of DEGs in *Setd1a*^+/-^ mice after TAK-418 treatment across NAc (top left), MD (top right), dStr (bottom left) and mPFC (bottom right). **b**, Percentages of rescued and partially rescued genes in the four brain regions. **c**, Dot plot showing GO terms enriched among the down-regulated DEGs that were rescued or partially rescued by TAK-418 treatment across the four brain regions. The −log_10_(*P* value) of GO terms are color-coded. Purple, nervous system development processes; Green, cellular organization; Blue, cellular energy metabolism. **d**, Heatmaps showing expression changes of DEGs in dStr of *Setd1a*^+/-^ mice and corresponding H3K4me3 enrichment around the transcriptional start sites (TSS) of those genes. **e**, Averaged H3K4me3 signal around the transcriptional start sites (TSS) of genes down-regulated in dStr neurons of *Setd1a*^+/−^ mice in WT, Het, and TAK-418 treated samples. **f**, Bar graph showing GO terms enriched among the down-regulated genes with H3K4me3 peaks in the dStr. **g**, Representative genes showing increased RNA expression and H3K4me3 promoter occupancy after TAK-418 treatment. **h**, Left, heat maps of averaged ΔF/F Ca^2^ responses in the dStr of *Setd1a^+/-^* (top) or TAK-418 treated mice (bottom) aligned to stimulus onset. Middle, averaged peri-stimulus traces of Ca^2^ signals of PS. Dashed line, event onset. Right, quantification of averaged peak ΔF/F values (dot plot). Two-tailed, paired *t*-test. n=4 mice. All RNA-seq and H3K4me3 CUT&RUN analysis were performed using two independent biological replicates.

Given the altered neuronal activity observed during PPI recording and the robust transcriptome rescue, we next focused on the dStr to determine whether the transcriptome rescue correlates with H3K4me3 restoration as previous study has shown that promoter H3K4me3 is tightly linked to gene expression^50^. To assess whether TAK-418 treatment restored H3K4me3 levels, we performed H3K4me3 CUT&RUN using dStr from WT, *Setd1a*^+/−^, and *Setd1a*^+/−^ with TAK-418 treatment (Extended Data Fig.8a). We found that TAK-418 treatment generally increased the H3K4me3 signals at the sites with reduced H3K4me3 in *Setd1a*^+/−^ mice (Extended Data Fig.8b,c). Notably, most downregulated genes with reduced H3K4me3 levels in *Setd1a*^+/−^ mice showed at least partial recovery of the H3K4me3 levels after the TAK-418 treatment (Fig.6d). At a global level, H3K4me3 enrichment at the transcription start sites (TSSs) of the down-regulated genes in the *Setd1a*^+/−^ mice was markedly decreased, but increased after TAK-418 treatment (Fig.6e). GO analysis of the rescued down-regulated genes with H3K4me3 peaks were enriched for cognitive function, synapse assembly and nervous system development (Fig.6f). For example, *Bsn*, which is critical for synapse assembly and maturation^51^, was down-regulated in *Setd1a*^+/−^ mice but rescued at both the RNA and H3K4me3 levels following the TAK-418 treatment (Fig.6g). Similar restoration was also observed for *Kmt2a*, *Nrxn2* and *Shank3*, genes essential for synaptic plasticity^52^, synapse assembly and cognition^53,54^ (Fig.6g; Extended Data Fig.8d). Furthermore, TAK-418 treatment significantly restored the neuronal response upon PS in PPI assay in *Setd1a*^+/−^ mice (Fig.6h), linking the restoration of neuronal activity dynamics to the rescue of SCZ-related behaviors.

Collectively, the above results demonstrate that TAK-418 treatment is able to reverses *Setd1a* heterozygosity-induced transcriptional, synaptic, and behavioral abnormalities through restoration of H3K4me3-mediated gene regulation. Broadly, our study provides mechanistic and therapeutic insights into how epigenetic modulation can correct transcriptional dysregulation underlying SCZ, establishing a framework for precision neuroepigenetic interventions in psychiatric disorders.

## Discussion

Great efforts in understanding the genetic cause of SCZ have resulted in the identification of genes whose mutation is a risk factor. Understanding how these genetic alterations contribute to SCZ pathogenesis is essential for developing effective treatment. In this study, we reveal that Setd1a haploinsufficiency confers brain region-specific vulnerability across multiple SCZ-relevant regions, linking epigenetic dysregulation to distinct transcriptional, neuronal, and behavioral abnormalities. By combining conditional knockout models (Fig.1), calcium imaging (Fig.2), transcriptomic profiling (Fig.3), and pharmacological rescue (Fig.4), we reveal that *Setd1a* loss-of-function disrupts neuronal activity and gene expression, impairs executive and motivational functions in the prefrontal cortex, sensorimotor gating in the striatum and thalamus. Importantly, we demonstrate that pharmacological inhibition of H3K4 demethylases, particularly LSD1, by TAK-418 effectively restores transcription, H3K4me3, and behavioral performance (Fig. 5 and 6). Our findings suggest that regional differences in epigenetics and transcription may underlie the heterogeneous symptoms of SCZ, raising the intriguing possibility that mechanistic-based epigenetic therapy may provide a promising treatment for SCZ.

### Region-specific vulnerability and systemic approaches in understanding schizophrenia

Unlike previous single-region or single-pathway models of SCZ, such as *Disc1*, *Shank3*, *Nrg1*, and *Grin2a* mutations, that revealed convergent yet spatially confined pathologies^55–58^, our study establishes *Setd1a* as a master regulator that orchestrates region-specific transcriptional, neuronal, and behavioral dysfunctions across multiple brain regions. By integrating epigenetic profiling, transcriptome, *in vivo* calcium recording, and conditional knockout behavioral screening, we delineate how heterozygosity of a single histone methyltransferase can reconfigure distinct molecular programs in cortical and subcortical regions, resulting in phenotypes characteristic of SCZ. Our findings suggest that SCZ could emerge from widespread yet regionally distinct network perturbations in contrast to the traditional uniform deficits localized to one brain structure^7,8^. Similar multi-omics approaches have been used to map transcriptomic and proteomic alterations across multiple brain regions in other SCZ-associated mutations, such as Grin2a^59^, *Srrm2*^60^, Sp4^61^ and *Zmym2*^61^. Such type of studies should help reveal the complex molecular logic of this disease.

The multi-dimensional, brain-wide approach of our study represents a departure from traditional approaches by bridging molecular alterations to systems level phenotypes, revealing that *Setd1a* deficiency triggers region-specific transcriptomic alterations that translate into behavioral dysfunctions through modulating neuronal activities. For instance, in the dStr, reduced H3K4me3 levels decreased expression of genes critical for nervous system development, synapse assembly, and cognitive function (Fig.3 and 6), leading to disrupted neuronal responses during sensorimotor gating (Fig.2) and PPI deficits (Fig.1). Remarkably, pharmacological restoration of H3K4me3 via demethylation inhibition largely rescued these deficits (Fig.5 and 6), demonstrating a causal and reversible link between epigenetic dysregulation and brain region dysfunction. In contrast, in the mPFC, *Setd1a* loss altered a different transcriptional network governing motivational and reward processing, resulting in anhedonia-like behavior. While loss of *Setd1a* in the NAc produced pronounced deficits in PPI, highlighting its contribution to sensorimotor gating and dopaminergic integration^62^, and *Setd1a* deletion in MD disrupted both PPI and social novelty recognition, consistent with the notion that the thalamic area is involved in coordination of social cognition and sensory filtering^63^. Although reward processing is frequently attributed to striatal circuits, particularly the dorsal striatum in action outcome learning and habit formation, accumulating evidence indicates that the mPFC plays an equally critical role in evaluating reward value, motivational drive, and behavioral flexibility^64^. The absence of sucrose preference impairments in striatal *Setd1a* cKO does not conflict with established striatal reward theories; instead, it highlights that *Setd1a* deficiency perturbs the valuation axis of reward processing selectively within the mPFC, whereas striatal dysfunction may engage more action contingent or reinforcement learning domains. This reinforces the broader theme of our study that heterozygous disruption of a single chromatin regulator yields anatomically dissociable phenotypes grounded in region-specific molecular logic. We also observed that Setd1a loss in the hippocampus alters social novelty preference. Given the established role of *Setd1a* in maintaining neural stem cell quiescence during adult hippocampal neurogenesis^65^, this finding raises the possibility that disrupted adult hippocampal neurogenesis may contribute to the observed behavioral phenotypes, an idea that needs further investigation.

By demonstrating that *Setd1a* acts as a region-specific epigenetic regulator that controls transcriptional precision within functionally distinct network, our study calls for the understanding of psychiatric pathogenesis from a dynamic, brain-wide systemic approach to reveal region-specific molecular perturbations and their link to neuronal and behavioral alterations. Consequently, therapeutic strategies that target a single brain region may fail to capture the systemic benefit achievable through global rebalancing of chromatin states, as demonstrated by the efficacy of H3K4 demethylase inhibition in restoring both transcriptional and behavioral integrity.

### Epigenetic rescue and its advantages to the traditional symptom relief treatment

Taking advantage of the fact that epigenetic modifications are reversible, we reasoned that loss of H3K4 methylation caused by *Setd1a* heterozygosity could be compensated by inhibition of H3K4 demethylation. Among the inhibitors tested, TAK-418 emerged as the most effective capable of rescuing H3K4me3 level, transcriptome, and behavioral phenotypes (Fig.4d and 5b). TAK-418 normalized synaptic gene expression, reinstated H3K4me3 occupancy at promoters of neuronal regulators (Fig.6g; Extended Data Fig.8d), and ameliorated deficits in behavior (Fig.5b). Compared with the KDM5 inhibitor JQKD82 or another LSD1 inhibitor ORY-1001, TAK-418 produced a broader restoration of the dysregulated genes and also rescued both the anhedonia and PPI phenotypes, which were not observed when using other inhibitors. By rescuing the gene expression, TAK-418 treatment addresses the root cause of the behavioral phenotypes, which is advantageous to the conventional symptomatic treatments for SCZ. Although *Setd1a* is mainly regarded as a H3K4me3 methyltransferase^16^, restoration of H3K4me1/2 via LSD1 inhibition may facilitate subsequent trimethylation by the remaining SETD1A in the *Setd1a*^+/−^ mice. In contrast, KDM5 inhibitors directly target H3K4me3 demethylation and risk excessive or nonspecific elevation of active chromatin marks, potentially disrupt transcriptional homeostasis. Further studies are needed to determine whether this potential mechanistic distinction is true.

Traditional antipsychotic therapies primarily target dopamine or glutamate signaling most notably through D /D receptor blockade^66,67^, which acutely modulates neurotransmission and alleviates certain positive symptoms. However, these treatments remain limited in efficacy and scope, as they fail to normalize the opposing dopaminergic and glutamatergic imbalances between the striatum and frontal cortex, and show minimal benefit for cognitive or negative symptoms. In contrast, epigenetic interventions modulate chromatin and reinstate physiological gene expression programs rather than transiently altering neurotransmitter flux. By restoring transcriptional networks across multiple brain regions and biological pathways simultaneously, such intervention addresses the root cause of the disease, thereby overcoming the restricted, symptom-specific effects of conventional single-pathway modulation. Notably, the fact that *Setd1a* loss perturbs several SCHEMA genes (Fig.3d) suggests that *SETD1A* heterozygosity disrupts a broader convergent molecular architecture shared across genetically diverse forms of SCZ. TAK-418 normalized expression of some of these dysregulated SCHEMA genes, indicating that LSD1 inhibition is not merely correcting Setd1a-specific downstream defects but may be restoring a wider transcriptional framework underpinning disease risk. This observation raises the possibility that epigenetic rebalancing through LSD1 inhibition could have therapeutic utility beyond SETD1A deficiency itself, potentially beneficial to other SCZ subtypes characterized by disruptions in chromatin-regulated synaptic and developmental pathways. Beyond SETD1A-related SCZ, the insights revealed in our study underscore a broader principle that psychiatric disorders may emerge from regionally distinct manifestations of a shared molecular vulnerability, and that targeting epigenetic regulators to restore gene expression fidelity may offer a promising strategy for treating other psychiatric disorders.

## Method

### Animals

All experiments were conducted in accordance with the National Institute of Health’s (NIH) *Guide for the Care and Use of Laboratory Animals* and approved by the Institutional Animal Care and Use Committee of Boston Children’s Hospital and Harvard Medical School. The *Setd1a* heterozygous (*Setd1a*^+/−^) and flox (*Setd1a*^flox/flox^) mice were generated in our lab. For molecular profiling, 8- to 14-week-old adult male mice were used. For behavioral assays, 12- to 16-week-old mice were used. The mice were housed in groups (3-5 mice per cage) in a 12 h light/dark cycle (light time, 07:00-19:00 h), with food and water provided ad libitum unless otherwise specified. Ambient temperature (23-25 °C) and humidity (55-62%) were automatically controlled.

*Setd1a*^flox/flox^ mice was generated by inserting loxP sites into the introns upstream of exons 14 and 17 using the CRISPR/Cas9 system, as previously described^68^ with minor modifications. Briefly, B6D2F1 (BDF1) mice were used for *in vitro* fertilization to obtain two-cell stage embryos. A mixture containing donor DNA harboring two loxP sites (30 ng/μl), Cas9 protein (20 ng/μl), and two sgRNAs (20 ng/μl each, sgRNA-1: 5’-GAT GGT GAG AAT ACT CAG TA-3’; sgRNA-2: 5’-CTA GAG GGT GAT AGG TAT TG-3’) was injected into two-cell embryos (20 hours post-fertilization) using a Piezo impact-driven micromanipulator (Primer Tech). The injected embryos were cultured in KSOM medium for 6 hours before being transferred into the oviducts of pseudo-pregnant Institute of Cancer Research (ICR) female mice (Charles River). F0 chimera mice were genotyped, and those carrying the correct loxP knock-in allele were backcrossed with wild-type C57BL/6J mice for two generations to establish the floxed line.

### Tissue dissection, RNA isolation, library preparation, and sequencing

Brain tissues for bulk RNA sequencing were prepared as preciously described^16^. To minimize biological variability, only male mice were used. *Setd1a*^+/−^ mice, their wild-type littermates and inhibitors treated mice were anesthetized by CO_2_ inhalation. Brains were rapidly dissected and rinsed in ice-cold phosphate-buffered saline (PBS). Then brains were cut into 1-mm coronal sections with a brain matrix, and the mPFC, NAc, dStr and MD were dissected and stored in TRIzol until further processing. The RNA was extracted from TRIzol with the Direct-zol RNA Microprep Kit (Zymo) following the manufacturer’s instruction, and the RNA from each sample was eluted in 10 μl of ribonuclease (RNase)-free water and stored at −80°C until further processing. The Smart-Seq V4 Kit (Takara) was used for preparing RNA-seq libraries. The RNA of each sample was used for cDNA generation following the manufacturer’s protocol with nine polymerase chain reaction (PCR) cycles to amplify the cDNA. After measuring cDNA concentration (Qubit dsDNA HS Assay Kit, Thermo Fisher Scientific) and quantifying control with Bioanalyzer 2100 (Bioanalyzer High Sensitivity DNA Analysis, Agilent), 150 ng of cDNA was used for library preparation with the Nextera XT DNA Library Preparation Kit (Illumina) according to the manufacturer’s instruction. The libraries were sequenced on Illumina NextSeq 1000 sequencer with pair-end sequencing (Read1, 76 bp; Read2, 76 bp).

### CUT&RUN

To collect tissues for CUT&RUN, each subregion was dissected the same as for RNA sequencing. Fresh tissues were homogenized in 1 ml of ice-cold homogenization buffer [320 mM sucrose, 5 mM CaCl_2_, 3 mM Mg(Ac)_2_, 10 mM tris (pH 7.6), 0.1 mM EDTA, 0.1% NP-40, 0.1 mM phenylmethylsulfonyl fluoride (PMSF), 1% BSA, and 1× protease inhibitor cocktails (Sigma-Aldrich)] using a 1-ml Dounce homogenizer (Wheaton). After 5 mins on ice, the homogenate was filtered with 40-μm cell strainer (Thermo Fisher Scientific) and diluted with 1 ml of dilution buffer [50% OptiPrep density gradient medium (Sigma-Aldrich), 5 mM CaCl_2_, 3 mM Mg(Ac)_2_, 10 mM tris (pH 7.6), 0.1 mM PMSF]. Lysate (0.5 ml) was loaded on the top of 0.5 ml of 29% iso-osmolar OptiPrep solution (in PBS) in a 1.5-ml centrifuge tube and centrifuged at 6000g for 10 min at 4°C. After removing the supernatant, the nuclei were resuspended in wash buffer [2.5 mM MgCl2, 1% BSA in PBS].

The H3K4me3 CUT&RUN libraries were prepared as previously described with some modifications^16^. Briefly, the extracted nuclei were then captured with BioMagPlus Concanavalin A (Polysciences) beads and incubated with a primary antibody (H3K4me3) for 16 hours at 4°C in antibody incubation buffer [20 mM Hepes (pH 7.5), 150 mM NaCl, 0.5 mM spermidine, 1× protease inhibitor cocktails (EDTA-free tablet, Sigma-Aldrich), 2 mM EDTA, and 0.005% digitonin (Life Technologies)]. The H3K4me3 antibody from Cell Signaling Technology (9727S; 1:100) was used. After unbound antibodies were washed away, protein A-MNase (micrococcal nuclease) (pA-MN; a gift from S. Henikoff) was added at a 1:280 ratio (500 ng/ml) and incubated for 3 hours at 4°C. After washing, CaCl_2_ was added to a final concentration of 2 mM to activate pA-MN, incubated for 20 min at 4°C, and then stopped by adding 1/10 volume of 10× STOP buffer [1700 mM NaCl, 100 mM EDTA, 20 mM EGTA, RNase A (250 μg/ml; Thermo Fisher Scientific), and glycogen (250 μg/ml;Sigma-Aldrich)]. The protein-DNA complexes were released by incubating at 37°C for 10 min, followed by 16,000*g* centrifugation for 5 min at 4°C. The supernatant was transferred to a new Lo-bound tube, 1/100 volume of 10% SDS, and 1/80 volume of Proteinase K (25 mg/ml; Thermo Fisher Scientific) were added and incubated at 55°C for at least 1 hour. DNA was then precipitated by phenol/chloroform/isoamylalcohol (25:24:1), followed by ethanol precipitation with glycogen, and then dissolved in water.

Sequencing libraries were prepared using the NEBNext Ultra II DNA library preparation kit for Illumina (New England Biolabs) according to the manufacturer’s instructions with a few modifications. Briefly, end repair was conducted at 20°C for 30 min, followed by dA-tailing at 65°C for 30 min. After adaptor ligation at 20°C for 30 min, the DNA fragments were purified by 1.8× volume of SPRIselect beads (Beckman Coulter) followed by 12 cycles (for H3K4me3) of PCR amplification with NEBNext Ultra II Q5 Master Mix (New England Biolabs). The PCR products were cleaned up with 0.9× volume of SPRIselect beads. The CUT&RUN libraries were quantified using the Qubit dsDNA HS Assay Kit (Agilent), and quality control was performed with the Bioanalyzer High Sensitivity DNA Analysis (Agilent). The libraries were sequenced on the Illumina NextSeq 1000 with pair-end 76-bp reads.

### FISH and imaging

Mice were transcardially perfused with PBS followed by 4% paraformaldehyde. Brains were cryoprotected in 30% sucrose for 2 days, embedded in OCT, and sectioned coronally at 16 μm (FISH) or 35 μm using cryostat (Leica CM3050 S). For FISH, sections were mounted on SuperFrost Plus slides, air-dried, and processed using the RNAscope Fluorescent Multiplex Assay (ACD Bioscience) with probes for *Setd1a* (cat. No. 551391-C1). For fluorescence imaging, sections were washed in PBS, followed by DAPI staining for 20 min at room temperature. Images were acquired using a Zeiss LSM800 confocal microscope with EC Plan-Neofluar 10×/0.30 M27 or Plan-Apochromat 20×/0.8 M27 objectives.

### Primary cortical neuronal cultures

Cortical neurons from P0 pups were dissected in Hanks’ balanced salt solution and digested with 0.25% trypsin at 37°C for 20 min and dissociated with a 1-ml pipette. Neurons were plated at the density of 1 × 10^5^/ml into Dulbecco’s modified eagle’s medium mixture F12 (Gibco) containing 10% fetal bovine serum on 6-well-plates with poly-D-lysine (50 μg/ml; Sigma-Aldrich). Neurons attached to the substrate were incubated with neurobasal-A medium (Gibco) containing B-27 (Gibco) supplements and L-GlutaMAX (Gibco). The cultures were maintained in a humidified atmosphere of 5% CO_2_ at 37°C. After 24 hours in culture, 10 mM cytosine-β-D-arabinofuranoside (Sigma-Aldrich) was added to restrict glial cell growth. Medium was half-changed every 2 days.

### FM dye imaging of presynaptic exocytosis in cultured neurons

Neurons cultured on glass-bottom dishes were imaged at DIV18. Prior to dye loading, cells were equilibrated for 10 min in Ca^2^ -free, low-K buffer (140 mM NaCl, 5 mM KCl, 5 mM NaHCO□, 1.2 mM NaH PO□, 1 mM MgCl□, 10 mM glucose, and 10 mM HEPES, pH 7.4). Neurons were then incubated for 5 min with 10 μM FM4-64 (Molecular Probes, Thermo Fisher Scientific) in high-K buffer (95 mM NaCl, 50 mM KCl, 1 mM MgCl□, 5 mM NaHCO□, 1.2 mM NaH PO□, 1.33 mM CaCl□, 10 mM glucose, and 10 mM HEPES, pH 7.4). Excess surface-bound dye was removed by washing cells for 10 min in Ca^2^ -free, low-K buffer.

Imaging was performed on a Zeiss LSM 800 confocal microscope equipped with a 63× oil-immersion objective (NA 1.4). FM4-64 fluorescence was excited using a 568-nm argon laser, and images were acquired at 512 × 512-pixel resolution. Following a 50 s baseline acquisition, neurons were depolarized with high-K buffer and imaged for an additional 350 s at 10-s intervals. Presynaptic puncta were selected for analysis, and fluorescence intensities were quantified from 1.5 × 1.5 μm regions of interest. All experiments were conducted at room temperature (25 °C). Image analysis was performed using Fiji.

### AAV vectors

The following AAV vectors used in this study were purchased from Addgene: pGP-AAV1-Syn-Flex-jGCaMP7s-WPRE (cat. no. 104491), pAAV-Ef1a-mCherry-IRES-Cre (cat. no. 55632), pAAV-Ef1a-mCherry (cat. no. 114470).

### Chemicals

ORY-1001 was purchased from Cayman Chemical (Cat# 19136). Cpd-48 and KDM5-C70 were gifts from Dr. Qin Yan lab (Yale School of Medicine). CPI-455 and JQKD82 were gifts from Dr. Jun Qi lab (Dana-Farber Cancer Institute). TAK-418 was a gift from Dr. Ed Scolnick in Stanley Center for Psychiatric Research, Broad Institute of MIT and Harvard.

### Stereotaxic brain surgeries

The injection was performed using a small-animal stereotaxic instrument (David Kopf Instruments, model 940) under general anesthesia by isoflurane (0.8 l min^−1^; isoflurane concentration 1.5%) in oxygen. A feedback heater was used to keep mice warm during surgeries. Once the mouse skull was exposed, a cranial window (1-2 mm^2^) was drilled unilaterally (for in vivo photometry, monosynaptic rabies and retrograde tracing experiments) or bilaterally (for optogenetic and chemogenetic experiments). Next, a glass capillary was lowered into the window to deliver approximately 0.2 μl of AAV vectors to the area of interest. The viral solution was delivered at a rate of 1 nl s^−1^ using a nanoliter injector (Nanoject III, Drummond Scientific 3000207). Following delivery, the pipette was left in place for 10 min and carefully withdrawn. For in vivo photometry experiments, following viral injection, a fiber optic cannula (100 μm in diameter; Inper Inc.) was implanted 0.1 mm above the viral injection site and secured with dental cement (Parkell, cat. no. S380). The coordinates of viral injection and implantation sites are based on previous literature and The Mouse Brain in Stereotaxic Coordinates (third edition) as follows: mPFC (anterior-posterior (AP), +2.10 mm; medial-lateral (ML), −0.4 mm; dorsal-ventral (DV), 1.8 mm); NAc (AP, +1.34 mm; ML, −0.53 mm; DV, 3.74 mm, with 10-degree angle); dStr (AP, +0.86 mm; ML, 1.8 mm; DV, 1.7 mm, with 10-degree angle) and MD (AP, −1.06 mm; ML, +0.5 mm; DV, −2.75 mm). Mice were allowed to recover in a warm blanket before they were transferred to housing cages for 2-4 weeks before behavioral evaluation was performed.

### Fiber photometry

GCaMP fluorescence was recorded using an RZ10X fiber photometry system (Tucker-Davis Technologies), which integrates LED drivers, excitation sources, and photosensors. Excitation light was delivered at 405 nm (isosbestic control) and 465 nm (GCaMP excitation). Emitted fluorescence was collected through a Mini Cube (Doric Lenses) and spectrally separated into two channels: 420-450 nm for autofluorescence and 500-550 nm for GCaMP7 signal detection. Mice were connected to an optical patch cable via an implanted cannula, and LED power at the fiber tip was adjusted to ∼20 μW to minimize photobleaching. Behavior was simultaneously recorded with CCD cameras (SuperCircuits).

In fiber photometry experiments, GCaMP fluorescence increased or decreased in response to neuronal excitation or inhibition, respectively. For pre-pulse inhibition (PPI) experiments, mice were habituated to the fiber optic cord for 10 min before recording. Behavioral events (acoustic startle [AS] or prepulse startle [PS] trials) were synchronized with the fluorescence signals using time-locked video recordings. For sucrose preference experiments, behavioral epochs (baseline exploration, initiation and termination of sucrose or water drinking) were manually annotated and synchronized with the fluorescence data.

Voltage signals corresponding to 405 nm and 465 nm excitation were acquired using Synapse software (Tucker-Davis Technologies) and exported for analysis as previously described^69^. Data were segmented by individual behavioral events. To calculate relative fluorescence changes (ΔF/F), the isosbestic (405 nm) signal was fitted to the GCaMP7 signal using a polynomial linear regression, and ΔF/F was computed as: ΔF/F = (GCaMP7 signal – fitted isosbestic) / fitted isosbestic. For calcium traces and heat maps, data were averaged within each mouse across multiple trials, with approximately 12 trials for AS and 8 trials for PS per mouse during the recording sessions. Each data point represents one mouse. Baseline fluorescence was defined as the mean ± SEM signal during the 5 s period preceding each stimulus. Peak responses were quantified within a 3 s window following event onset, with time 0 s indicating the start of the event.

### Behavioral assays

Behavioral measurements were conducted sequentially as below, with a minimum of one-day interval between tests to minimize stress-induced confounds. All behavioral tests were performed consistently in the afternoon to maintain uniformity across experimental batches.

#### Open field tests

A clear box (27.3 cm × 27.3 cm square base with 20.3 cm high walls) was used for the open field test, and the center zone was 36% of the total area. Before testing, mice were habituated to the test room for at least 20 min. Mice were placed in the center of the box at the start of the assay. Movements were recorded (Med Associates, ENV-510) for 1 h in 5 min bins. In addition to regular parameters related to locomotor activity (such as total travel distance, velocity, ambulatory time, resting time), time spent and distance traveled in the center area of the testing arena were also recorded and analyzed.

#### Y-maze

Mice were allowed to adapt in a separate room for approximately 30 min before the test. Each mouse was run twice in the Y-maze: first, a 3-min habituation phase and then a 3-min test phase. The delay or intertrial interval (ITI) between the end of the habituation phase and start of the test phase is 2 min. During the habituation phase, one arm (either left or right) is blocked. The start arm always remains the same. At the end of the habituation phase, the mouse was placed back into the holding cage for the 2-min ITI. The blockade was then removed, and the maze was lightly cleaned. Then, the mouse was placed back into the start arm for the test trial and returned to the home cage after the trial is completed. Distance and time traveled in the maze were recorded by Noldus EthoVision XT.

#### Three-chamber social interaction

Each social interaction chamber (30 × 30 × 30 cm) contained dividing walls with an open middle section to allow access. Both outer chambers contained wire cups. Mice were given free access to the apparatus for 10 min (in the absence of other mice) to habituate and confirm initial unbiased preference. The time spent in each chamber was recorded, and the time spent in close interaction with the nose point within 2 cm of the enclosure was also recorded (EthoVision XT 14, Noldus Information Technology). To test for sociability, mice were placed into the middle chamber of the apparatus with one outer chamber containing one mouse (’stranger 1’) confined in a wire cup and the other chamber containing a Lego block. For social novelty preference, mice were again placed into the middle chamber with one chamber containing the familiar mouse (stranger 1) and the other containing a novel mouse (’stranger 2’) confined in a wire cup. Male familiar and novel mice introduced for assay in social interactions matched the male test subject. For each phase, the test mice explored the entire arena throughout the 10-min trial. The time spent interacting with the empty wire, stranger 1 and stranger 2 mice during the 10-min session was recorded.

#### Sucrose preference test

Mice were singly housed for at least 2 days before the tests. For the first 2 days, animals were habituated with two bottles of water, with the position of the two bottles switched at 24 h. On the night of day 2, mice were water-deprived for 16 h and then the test was started on day 3. The test period lasted for 3 h, during which the mice were exposed to one bottle of pure drinking water and one bottle of drinking water containing 2% sucrose solution (w:v). Bottles were weighed before and after the tests to measure the water consumption. The sucrose preference ratio was calculated by dividing the consumption of sucrose solution by the total consumption of both pure drinking water and sucrose solution.

#### Acoustic startle and PPI

Animals are submitted to sessions consisting of 10 blocks of 11 trials each (110 trials total). Within each block, various white noise acoustic stimuli (20 to 120 dB) are presented in a random order with a variable ITI of mean 15 s (10 to 20 s). The duration of the stimulus is 40 ms. Responses are recorded for 150 ms from startle onset and are sampled every millisecond. The order of stimuli is random. Habituation time is 5 min. No background white noise was applied. Before the testing, animals were placed in the PPI chambers for a 5-min session of white noise (70 dB) habituation. After this adaptation period, the test session was automatically started. The session began with a habituation block of six presentations of the startle stimulus alone, followed by 10 PPI blocks of six different types of trials. Trial types are null (no stimuli), startle (120 dB), startle plus pre-pulse (65, 75, or 85 dB, 5000-Hz tone) and pre-pulse alone (85 dB, 5000-Hz tone). Each PPI trial began with a 50-ms null period during which baseline movements were recorded. There was a subsequent 20-ms period during which pre-pulse stimuli were presented and responses to the pre-pulse measured. After further 100 ms, the startle stimuli were presented for 40 ms, and responses were recorded for 140 ms from startle onset. Responses were sampled every millisecond. The ITI is variable with an average of 15 s (range from 10 to 20 s). All PPI enclosures were cleaned with water and 70% ethanol following each test. The percent PPI was calculated as (100 – pre-pulse + startle/startle × 100) which provides a measure of sensorimotor gating performance.

### Statistics

All statistical analyses were performed using GraphPad Prism (version 10) software and fiber photometry results were analyzed by MATLAB. No statistical methods were used to pre-determine sample sizes but our sample sizes are similar to those reported in previous publications^69,70^. All mice were randomly assigned to different groups and data collection was randomized whenever possible. Mice that, after histological inspection, had the location of the viral injection (reporter protein), cannula implantation, or the optic fiber(s) outside the area of interest were excluded. Data collection and analysis were not performed blind to the conditions of the experiments. Most behavioral experiments were controlled by an automated computer system, and the data were collected and analyzed in an unbiased way. Statistical analyses were two-tailed. Parametric tests including paired and unpaired *t*-test and one-way ANOVA were used if distributions passed the Kolmogorov-Smirnov normality test. Normality tests were not performed for one-way ANOVA with missing values. If data were not normally distributed, non-parametric tests were used. One-sample *t*-test was performed to determine whether the group mean differed from a specific value. For comparisons across more than two groups, one-way ANOVA or repeated-measures one-way ANOVA was performed for normally distributed data, followed by Tukey’s multiple comparison tests; if the SDs are significant different, Welch ANOVA test with Dunnett’s T3 multiple comparisons test was performed instead; two-way ANOVA was performed for differences between groups with two independent variables, followed by Šídák’s multiple comparison tests. For detailed statistical analysis, see the figure legends.

### RNA-seq data and GO enrichment analysis

Raw sequencing reads were trimmed using Trimmomatic^71^ (v0.39) to remove sequencing adaptors, and subsequently mapped to the GRCm38 genome using STAR^72^ (v2.7.8a). Gene expression was quantified with featureCounts^73^ (v2.0.1) by counting reads mapped to each gene. Then, edgeR^74^ (v3.32.1) was employed for normalization and differential expression analysis. Genes with RPKM lower than 1 were defined as lowly expressed genes and excluded from the differential expression analysis. Differentially expressed genes (DEGs) were identified using quasi-likelihood F-test (glmQLFit and glmQLFTest functions from edgeR). Genes with a *p*-value below 0.01 and an absolute value of fold change greater than 1.2 were defined as DEGs. Gene Ontology (GO) enrichment was performed using the R package clusterProfiler^75^. Gene Set Enrichment Analysis (GSEA) was performed using clusterProfiler^75^ and enrichplot based on gene list from Published paper^76,77^.

### CUT&RUN data analysis

Raw sequencing reads were trimmed using Trimmomatic^71^ (v0.39) to remove sequencing adaptors, and subsequently aligned to the GRCm38 reference genome using bowtie2^78^ (v2.4.2) with parameters: --local --very-sensitive-local --no-unal --no-mixed --no-discordant --dovetail -I 10 -X 700 --soft-clipped-unmapped-tlen. PCR duplicates were removed by Picard MarkDuplicates (v2.23.4) and reads with a mapping quality below 30 were removed. Then, MACS2^79^ (v2.2.7.1) was used to call significant peaks with parameters "-f BAMPE -B --SPMR - q 0.05 -g mm --keep-dup all". To obtain highly conserved H3K4me3 binding sites, only peaks present in both biological replicates were called as binding sites. Normalization scale factors were first estimated using csaw (v1.40.0) ^80^, and the signal tracks were generated with deeptools^81^ bamCoverage (v3.5.1) with bin size of 1 and normalized using CPM. The heatmaps of binding profiles were calculated with deeptools^81^ computeMatrix (v3.5.1) using normalized bigwig signal tracks as input and bin size of 10 and visualized in R with profileplyr and EnrichedHeatmap^82^ packages.

### Transcription factor enrichment analysis

For transcription factor (TF) enrichment analysis, the enrichment of up- and downregulated DEGs was evaluated for each brain region using the ChEA3^41^ overrepresentation test. TFs of interest were defined as those that were differentially expressed in *Setd1a*^+/−^ mutants within the brain region of interest and were associated with *Setd1a* in the ChEA3 database. For each selected TF, the proportion of DEGs associated with that TF was calculated by counting the number of DEGs linked to it in any of the ChEA3 libraries and dividing this count by the total number of DEGs tested in the enrichment analysis.

## Data and code availability

All data are available in the main text or supplementary materials. Sequencing data that support the findings of this study have been deposited in the Gene Expression Omnibus (GEO) under accession code GSE313686.

## Materials availability

Related reagents and mouse lines described in this study are available after signing standard Biological Material Transfer Agreement.

## Acknowledgement

We thank the Mouse Behavior Core of Harvard Medical School and its director Dr. Barbara Caldarone for her help; Dr. Kris Dickson of the Stanley Center for Psychiatric Research at the Broad Institute of MIT and Harvard for critical reading of the manuscript, and Dr. Qianying Yang of the Zhang lab for technical assistance. This project was partly supported by a grant from the Stanley Center for Psychiatric Research at the Broad Institute. Y.L. was supported by the Tommy Fuss Fund. Y.Z. is an investigator of the Howard Hughes Medical Institute.

## Author contributions

Y.Z. conceived the project; Y.L. and Y.Z. designed the experiments; Y.L. and G.X. performed behavioral tests, inhibitors screen and fiber photometry recording. Y.L. generated all sequencing libraries with S.J.’s help. C.J.Z. generated *Setd1a*^flox/flox^ mouse line. G.X. performed fiber photometry analysis. S.J. analyzed the sequencing data and C.Z. performed the preliminary analysis. J.Q. provided JQKD82. E.S. and M.S. participated discussion and made helpful suggestions. Y.L., G.X. S.J. and Y.Z. interpreted the data. Y.L. and Y.Z. wrote the manuscript with inputs from all authors.

## Competing interests

The authors declare no competing interests.

## Supplemental Figure Legends

**Extended Data Fig.1 Brain activation following acoustic startle response and generation of the *Setd1a*^flox/flox^ mouse line.**

**a**, Representative images showing c-Fos expression in control and acoustic startle response (ASR) groups. Scale bar, 100 μm.

**b**, Quantification of the average number of c-Fos+ neurons per 0.5 mm^2^ in the medial prefrontal cortex (mPFC), medial nucleus accumbens (NAcMed), dorsal striatum (dStr) and mediodorsal thalamus (MD). *n* = 3 sections from three mice of each group; Two-tailed, unpaired t tests.

**c**, Schematic of the gene-targeting strategy used to generate the *Setd1a*^flox/flox^ mouse line.

**d**, Genotyping results obtained by polymerase chain reaction (PCR).

Data were presented as means ± SEM. ns, not significant; \*\**P* < 0.01, \*\*\**P* < 0.001.

**Extended Data Fig.2 Behavioral features of *Setd1a* cKO mice**

**a**, Behavioral screen results of open field and three-chamber social interaction in WT and mPFC *Setd1a* cKO mice.

**b**, Behavioral screen results of open field, Y maze and three-chamber social interaction test in WT and NAc *Setd1a* cKO mice.

**c**, Behavioral screen results of open field, Y maze and three-chamber social interaction test in WT and dStr *Setd1a* cKO mice.

**d**, Behavioral screen results of open field, Y maze and three-chamber social interaction test in WT and MD *Setd1a* cKO mice.

**e**, Behavioral screen results of open field, Y maze and three-chamber social interaction test in WT and HPC *Setd1a* cKO mice.

**f**, Behavioral screen results of open field, Y maze and three-chamber social interaction test in WT and cerebellum (CB) *Setd1a* cKO mice.

Data were presented as means ± SEM. Each dot represents the result from an individual mouse. Two-tailed unpaired Student’s *t* test. ns, not significant; \**P* < 0.05, \*\**P* < 0.01, \*\*\**P* < 0.001, \*\*\*\**P* < 0.0001.

**Extended Data Fig.3 Setd1a loss induces brain region–specific transcriptome alterations**

**a-d**, Correlations of RNA-seq replicates from each brain region. cor, Pearson correlation coefficient. The x and y axis of the dot plots are Log (CPM+1).

**e**, Dot plot showing the Gene Ontology (GO) terms enriched for the up-regulated DEGs of four different brain regions. The −log_10_(*P*) of GO terms are color-coded.

**f**, Bar plot showing the representative genes expression across brain regions. Two-tailed unpaired t tests.

**g**, TF enrichment analysis results showing target genes of the denoted TFs and their percentage overlap between up-regulated DEGs in NAc, MD and dStr.

Data were presented as means ± SEM. Each dot represents the result from one replicate. ns, not significant; \**P* < 0.05, \*\**P* < 0.01, \*\*\**P* < 0.001.

**Extended Data Fig.4 Effects of H3K4 methylation inhibitors on neurite outgrowth *in vitro***

**a**, Quantification of normalized neurite numbers grow in vitro for 10 days of WT, *Setd1a*^+/−^, and *Setd1a*^+/−^ treated with different concentrations of ORY-1001.

**b-d**, Same as **a**, but treated with Cpd48 (**b**), Cpi455 (**c**) or KDM70 (**d**). n= 50 cells per group.

**Extended Data Fig.5 Transcriptomic profiling reveals reversal of *Setd1a*^+/-^ associated gene expression by H3K4 demethylase inhibitors**

**a**, Pearson correlation showing correlations among RNA-seq replicates from each experimental group. The x and y axis of the dot plots are Log (CPM+1).

**b**, Numbers of differentially expressed genes (DEGs) identified in each group; blue and red bars indicate down- and up-regulated DEGs, respectively.

**c**, Venn diagram of DEGs showing very little overlap between *Setd1a*^+/-^ affected genes and ORY-1001 treatment affected genes. Left, down-regulated in *Setd1a*^+/-^ vs. up-regulated with ORY-1001; Right, up-regulated in *Setd1a*^+/-^ vs. down-regulated with ORY-1001.

**d-e**, GO enrichment analysis of DEGs following TAK-418 (**d**) and JQKD82 (**e**) treatment. GO terms are color-coded by –log (*p*) value.

**Extended Data Fig.6 Physiological and behavioral assessments following TAK-418 and JQKD82 treatment**

Behavioral assays following 3 weeks of JQKD82 treatment. Top, experimental timeline. Bottom, sucrose preference (left) and PPI (right) tests of WT, *Setd1a*^+/−^ and JQKD82 treated *Setd1a*^+/−^ mice. One-way ANOVA test with Tukey’s multiple comparisons test. n=8 per each group for sucrose preference test; n=10 for WT, n=8 for Het groups, n=10 for JQKD82 group for PPI test.

Each dot represents the result from an individual mouse.

Data were presented as means ± SEM. n.s., not significant; \**P* < 0.05, \*\**P* < 0.01, \*\*\**P* < 0.001.

**Extended Data Fig.7 TAK-418 treatment restores transcriptional alterations**

**a**, Pearson correlation showing the consistency among RNA-seq replicates across experimental groups. The x and y axis of the dot plots are Log (CPM+1).

**b**, Dot plot showing GO terms enriched among the rescued up-regulated DEGs following TAK-418 treatment across the four brain regions. The –log (*p*) of GO terms are color coded.

**Extended Data Fig.8 TAK-418 treatment restores H3K4m3 occupancy**

**a**, Pearson correlation showing the consistency among H3K4me3 CUT&RUN replicates across experimental groups. The x and y axis of the dot plots are log_2_(normalized_counts+1). Each dot represents a 5-kb bin.

**b**, Venn diagram showing the overlap of H3K4me3 peaks in WT or *Setd1a*^+/-^ dStr tissues.

**c**, Heatmap showing the *Setd1a*^+/-^ dStr loss H3K4me3 peaks that are restored or partially restored by TAK-418 treatment. The H3K4me3 signal is color-coded csaw normalized CPM. C represents peak center.

**d**, Representative genes showing increased RNA expression and H3K4me3 after TAK-418 treatment.

## Supplementary Tables

**Supplementary Table 1**: List of genes altered by Setd1a heterozygosity and rescued by H3K4 demethylase inhibitor treatment in different brain regions

**Supplementary Table 2**: List of genes altered by Setd1a heterozygosity and rescued by H3K4 demethylase inhibitor treatment of cultured cortical neurons

**Supplementary Table 3**: zScore of differentially expressed genes in dStr tissues of WT, *Setd1a^+/-^*, and *Setd1a^+/-^* mice treated with TAK-418 and the H3K4me3 binding status

**Supplementary Table 4**: Summary of sequenced libraries in this study

## Notes

### Competing Interest Statement

The authors have declared no competing interest.

## Reference

1 Kahn, R. S. et al. Schizophrenia. Nature Reviews Disease Primers 1, 15067, doi:10.1038/nrdp.2015.67 (2015).

2 Marder, S. R. & Cannon, T. D. Schizophrenia. N Engl J Med 381, 1753–1761, doi:10.1056/NEJMra1808803 (2019).

3 Smeland, O. B., Frei, O., Dale, A. M. & Andreassen, O. A. The polygenic architecture of schizophrenia — rethinking pathogenesis and nosology. Nature Reviews Neurology 16, 366–379, doi:10.1038/s41582-020-0364-0 (2020).

4 Stroup, T. S. & Gray, N. Management of common adverse effects of antipsychotic medications. World Psychiatry 17, 341–356, doi:10.1002/wps.20567 (2018).

5 Goff, D. C. The Pharmacologic Treatment of Schizophrenia-2021. Jama 325, 175–176, doi:10.1001/jama.2020.19048 (2021).

6 Young, J. W. & Geyer, M. A. Developing treatments for cognitive deficits in schizophrenia: the challenge of translation. J Psychopharmacol 29, 178–196, doi:10.1177/0269881114555252 (2015).

7 Wang, Y. et al. Symptom-circuit mappings of the schizophrenia connectome. Psychiatry Res 323, 115122, doi:10.1016/j.psychres.2023.115122 (2023).

8 Stephan, K. E., Baldeweg, T. & Friston, K. J. Synaptic plasticity and dysconnection in schizophrenia. Biol Psychiatry 59, 929–939, doi:10.1016/j.biopsych.2005.10.005 (2006).

9 Singh, T. et al. Rare coding variants in ten genes confer substantial risk for schizophrenia. Nature 604, 509–516, doi:10.1038/s41586-022-04556-w (2022).

10 Wang, S. et al. Loss-of-function variants in the schizophrenia risk gene SETD1A alter neuronal network activity in human neurons through the cAMP/PKA pathway. Cell Rep 39, 110790, doi:10.1016/j.celrep.2022.110790 (2022).

11 Mukai, J. et al. Recapitulation and Reversal of Schizophrenia-Related Phenotypes in Setd1a-Deficient Mice. Neuron 104, 471–487 e412, doi:10.1016/j.neuron.2019.09.014 (2019).

12 Nagahama, K. et al. Setd1a Insufficiency in Mice Attenuates Excitatory Synaptic Function and Recapitulates Schizophrenia-Related Behavioral Abnormalities. Cell Rep 32, 108126, doi:10.1016/j.celrep.2020.108126 (2020).

13 Chen, R. et al. Decoding molecular and cellular heterogeneity of mouse nucleus accumbens. Nature neuroscience 24, 1757–1771, doi:10.1038/s41593-021-00938-x (2021).

14 Su, X. et al. Mutations of schizophrenia risk gene SETD1A dysregulate synaptic function in human neurons. Mol Psychiatry 30, 5680–5693, doi:10.1038/s41380-025-03246-z (2025).

15 Takata, A., Ionita-Laza, I., Gogos, Joseph A., Xu, B. & Karayiorgou, M. De Novo Synonymous Mutations in Regulatory Elements Contribute to the Genetic Etiology of Autism and Schizophrenia. Neuron 89, 940–947, 10.1016/j.neuron.2016.02.024 (2016).

16 Chen, R. et al. Cell type-specific mechanism of Setd1a heterozygosity in schizophrenia pathogenesis. Sci Adv 8, eabm1077, doi:10.1126/sciadv.abm1077 (2022).

17 Hoshii, T. et al. SETD1A regulates transcriptional pause release of heme biosynthesis genes in leukemia. Cell Reports 41, doi:10.1016/j.celrep.2022.111727 (2022).

18 Shilatifard, A. The COMPASS family of histone H3K4 methylases: mechanisms of regulation in development and disease pathogenesis. Annu Rev Biochem 81, 65–95, doi:10.1146/annurev-biochem-051710-134100 (2012).

19 Pekowska, A. et al. H3K4 tri-methylation provides an epigenetic signature of active enhancers. Embo j 30, 4198–4210, doi:10.1038/emboj.2011.295 (2011).

20 Yao, B. et al. Epigenetic mechanisms in neurogenesis. Nature reviews. Neuroscience 17, 537–549, doi:10.1038/nrn.2016.70 (2016).

21 Park, J., Lee, K., Kim, K. & Yi, S.-J. The role of histone modifications: from neurodevelopment to neurodiseases. Signal Transduction and Targeted Therapy 7, 217, doi:10.1038/s41392-022-01078-9 (2022).

22 Ma, D. K. et al. Epigenetic choreographers of neurogenesis in the adult mammalian brain. Nature neuroscience 13, 1338–1344, doi:10.1038/nn.2672 (2010).

23 Smigielski, L., Jagannath, V., Rössler, W., Walitza, S. & Grünblatt, E. Epigenetic mechanisms in schizophrenia and other psychotic disorders: a systematic review of empirical human findings. Molecular Psychiatry 25, 1718–1748, doi:10.1038/s41380-019-0601-3 (2020).

24 Klengel, T. & Binder, Elisabeth B. Epigenetics of Stress-Related Psychiatric Disorders and Gene × Environment Interactions. Neuron 86, 1343–1357, 10.1016/j.neuron.2015.05.036 (2015).

25 Ludwig, B. & Dwivedi, Y. Dissecting bipolar disorder complexity through epigenomic approach. Molecular Psychiatry 21, 1490–1498, doi:10.1038/mp.2016.123 (2016).

26 Zhang, T., Cooper, S. & Brockdorff, N. The interplay of histone modifications – writers that read. EMBO reports 16, 1467–1481-1481, 10.15252/embr.201540945 (2015).

27 Bannister, A. J. & Kouzarides, T. Regulation of chromatin by histone modifications. Cell Res 21, 381–395, doi:10.1038/cr.2011.22 (2011).

28 Schroeder, F. A., Lin, C. L., Crusio, W. E. & Akbarian, S. Antidepressant-like effects of the histone deacetylase inhibitor, sodium butyrate, in the mouse. Biol Psychiatry 62, 55–64, doi:10.1016/j.biopsych.2006.06.036 (2007).

29 Baba, R. et al. Investigating the Therapeutic Potential of LSD1 Enzyme Activity-Specific Inhibition by TAK-418 for Social and Memory Deficits in Rodent Disease Models. ACS Chemical Neuroscience 13, 313–321, doi:10.1021/acschemneuro.1c00713 (2022).

30 Veerasakul, S., Thanoi, S., Reynolds, G. P. & Nudmamud-Thanoi, S. Effect of Methamphetamine Exposure on Expression of Calcium Binding Proteins in Rat Frontal Cortex and Hippocampus. Neurotox Res 30, 427–433, doi:10.1007/s12640-016-9628-2 (2016).

31 Lesh, T. A., Niendam, T. A., Minzenberg, M. J. & Carter, C. S. Cognitive control deficits in schizophrenia: mechanisms and meaning. Neuropsychopharmacology 36, 316–338, doi:10.1038/npp.2010.156 (2011).

32 McCutcheon, R., Beck, K., Jauhar, S. & Howes, O. D. Defining the Locus of Dopaminergic Dysfunction in Schizophrenia: A Meta-analysis and Test of the Mesolimbic Hypothesis. Schizophr Bull 44, 1301–1311, doi:10.1093/schbul/sbx180 (2018).

33 Baldan Ramsey, L. C., Xu, M., Wood, N. & Pittenger, C. Lesions of the dorsomedial striatum disrupt prepulse inhibition. Neuroscience 180, 222–228, doi:10.1016/j.neuroscience.2011.01.041 (2011).

34 Clinton, S. M. & Meador-Woodruff, J. H. Thalamic dysfunction in schizophrenia: neurochemical, neuropathological, and in vivo imaging abnormalities. Schizophr Res 69, 237–253, doi:10.1016/j.schres.2003.09.017 (2004).

35 Bledau, A. S. et al. The H3K4 methyltransferase Setd1a is first required at the epiblast stage, whereas Setd1b becomes essential after gastrulation. Development 141, 1022–1035, doi:10.1242/dev.098152 (2014).

36 Braff, D. et al. Prestimulus effects on human startle reflex in normals and schizophrenics. Psychophysiology 15, 339–343, doi:10.1111/j.1469-8986.1978.tb01390.x (1978).

37 Swerdlow, N. R., Geyer, M. A. & Braff, D. L. Neural circuit regulation of prepulse inhibition of startle in the rat: current knowledge and future challenges. Psychopharmacology (Berl) 156, 194–215, doi:10.1007/s002130100799 (2001).

38 Gandal, M. J. et al. Transcriptome-wide isoform-level dysregulation in ASD, schizophrenia, and bipolar disorder. Science 362, doi:10.1126/science.aat8127 (2018).

39 Ruzicka, W. B. et al. Single-cell multi-cohort dissection of the schizophrenia transcriptome. Science 384, eadg5136, doi:10.1126/science.adg5136 (2024).

40 Koopmans, F. et al. SynGO: An Evidence-Based, Expert-Curated Knowledge Base for the Synapse. Neuron 103, 217–234.e214, 10.1016/j.neuron.2019.05.002 (2019).

41 Keenan, A. B. et al. ChEA3: transcription factor enrichment analysis by orthogonal omics integration. Nucleic Acids Res 47, W212–W224, doi:10.1093/nar/gkz446 (2019).

42 Duclot, F. & Kabbaj, M. The Role of Early Growth Response 1 (EGR1) in Brain Plasticity and Neuropsychiatric Disorders. Front Behav Neurosci 11, 35, doi:10.3389/fnbeh.2017.00035 (2017).

43 Huang, X., Powell-Coffman, J. A. & Jin, Y. The AHR-1 aryl hydrocarbon receptor and its co-factor the AHA-1 aryl hydrocarbon receptor nuclear translocator specify GABAergic neuron cell fate in C. elegans. Development 131, 819–828, doi:10.1242/dev.00959 (2004).

44 Rottkamp, C. A., Lobur, K. J., Wladyka, C. L., Lucky, A. K. & O’Gorman, S. Pbx3 is required for normal locomotion and dorsal horn development. Developmental Biology 314, 23–39, 10.1016/j.ydbio.2007.10.046 (2008).

45 Mok, C. H. et al. PBX1 and PBX3 transcription factors regulate SHH expression in the Frontonasal Ectodermal Zone through complementary mechanisms. PLOS Genetics 21 (2025).

46 Kramer, J. M. & van Bokhoven, H. Genetic and epigenetic defects in mental retardation. The International Journal of Biochemistry & Cell Biology 41, 96–107, 10.1016/j.biocel.2008.08.009 (2009).

47 Wolf, A., Yitzhaky, A. & Hertzberg, L. SMAD genes are up-regulated in brain and blood samples of individuals with schizophrenia. J Neurosci Res 101, 1224–1235, doi:10.1002/jnr.25188 (2023).

48 Wu, L. et al. KDM5 histone demethylases repress immune response via suppression of STING. PLoS Biol 16, e2006134, doi:10.1371/journal.pbio.2006134 (2018).

49 Yin, W. et al. Safety, pharmacokinetics and pharmacodynamics of TAK-418, a novel inhibitor of the epigenetic modulator lysine-specific demethylase 1A. Br J Clin Pharmacol 87, 4756–4768, doi:10.1111/bcp.14912 (2021).

50 Barski, A. et al. High-Resolution Profiling of Histone Methylations in the Human Genome. Cell 129, 823–837, 10.1016/j.cell.2007.05.009 (2007).

51 Fu, J. M. et al. Rare coding variation provides insight into the genetic architecture and phenotypic context of autism. Nat Genet 54, 1320–1331, doi:10.1038/s41588-022-01104-0 (2022).

52 Jakovcevski, M. et al. Neuronal Kmt2a/Mll1 histone methyltransferase is essential for prefrontal synaptic plasticity and working memory. The Journal of neuroscience : the official journal of the Society for Neuroscience 35, 5097–5108, doi:10.1523/jneurosci.3004-14.2015 (2015).

53 Hallermann, S. et al. Bassoon Speeds Vesicle Reloading at a Central Excitatory Synapse. Neuron 68, 710–723, doi:10.1016/j.neuron.2010.10.026 (2010).

54 Hu, C., Chen, W., Myers, S. J., Yuan, H. & Traynelis, S. F. Human GRIN2B variants in neurodevelopmental disorders. Journal of Pharmacological Sciences 132, 115–121, 10.1016/j.jphs.2016.10.002 (2016).

55 Yin, D. M. et al. Reversal of behavioral deficits and synaptic dysfunction in mice overexpressing neuregulin 1. Neuron 78, 644–657, doi:10.1016/j.neuron.2013.03.028 (2013).

56 Vullhorst, D. et al. Selective Expression of ErbB4 in Interneurons, But Not Pyramidal Cells, of the Rodent Hippocampus. The Journal of Neuroscience 29, 12255, doi:10.1523/JNEUROSCI.2454-09.2009 (2009).

57 Alvarez, R. J., Pafundo, D. E., Zold, C. L. & Belforte, J. E. Interneuron NMDA Receptor Ablation Induces Hippocampus-Prefrontal Cortex Functional Hypoconnectivity after Adolescence in a Mouse Model of Schizophrenia. The Journal of Neuroscience 40, 3304, doi:10.1523/JNEUROSCI.1897-19.2020 (2020).

58 Zhou, T. et al. Enhancement of mediodorsal thalamus rescues aberrant belief dynamics in a mouse model with schizophrenia-associated mutation. bioRxiv, doi:10.1101/2024.01.08.574745 (2024).

59 Farsi, Z. et al. Brain-region-specific changes in neurons and glia and dysregulation of dopamine signaling in Grin2a mutant mice. Neuron 111, 3378–3396.e3379, 10.1016/j.neuron.2023.08.004 (2023).

60 Aryal, S. et al. Reduction of SynGAP-γ, disrupted splicing of <em>Agap3</em>, and oligodendrocyte deficits in <em>Srrm2</em> mice, a genetic model of schizophrenia and neurodevelopmental disorder. bioRxiv, 2024.2010.2010.617460, doi:10.1101/2024.10.10.617460 (2024).

61 Kwon, M. J. et al. Muti-omics characterization reveals brain-wide disruption of synapses and region- and age-specific changes in neurons and glia in <em>Sp4</em> mutant mice, a genetic model of schizophrenia and bipolar disorder. bioRxiv, 2024.2010.2012.618006, doi:10.1101/2024.10.12.618006 (2024).

62 Mansbach, R. S., Geyer, M. A. & Braff, D. L. Dopaminergic stimulation disrupts sensorimotor gating in the rat. Psychopharmacology (Berl) 94, 507–514, doi:10.1007/bf00212846 (1988).

63 Ferguson, B. R. & Gao, W. J. Thalamic Control of Cognition and Social Behavior Via Regulation of Gamma-Aminobutyric Acidergic Signaling and Excitation/Inhibition Balance in the Medial Prefrontal Cortex. Biol Psychiatry 83, 657–669, doi:10.1016/j.biopsych.2017.11.033 (2018).

64 Pastor, V. & Medina, J. H. Medial prefrontal cortical control of reward- and aversion-based behavioral output: Bottom-up modulation. Eur J Neurosci 53, 3039–3062, doi:10.1111/ejn.15168 (2021).

65 Zhao, T. et al. Epigenetic maintenance of adult neural stem cell quiescence in the mouse hippocampus via Setd1a. Nature Communications 15, 5674, doi:10.1038/s41467-024-50010-y (2024).

66 Kaar, S. J., Natesan, S., McCutcheon, R. & Howes, O. D. Antipsychotics: Mechanisms underlying clinical response and side-effects and novel treatment approaches based on pathophysiology. Neuropharmacology 172, 107704, doi:10.1016/j.neuropharm.2019.107704 (2020).

67 Gross, G., Wicke, K. & Drescher, K. U. Dopamine D receptor antagonism--still a therapeutic option for the treatment of schizophrenia. Naunyn Schmiedebergs Arch Pharmacol 386, 155–166, doi:10.1007/s00210-012-0806-3 (2013).

68 Zhou, C., Wang, M., Zhang, C. & Zhang, Y. The transcription factor GABPA is a master regulator of naive pluripotency. Nat Cell Biol 27, 48–58, doi:10.1038/s41556-024-01554-0 (2025).

69 Liu, Y. et al. A subset of dopamine receptor-expressing neurons in the nucleus accumbens controls feeding and energy homeostasis. Nat Metab 6, 1616–1631, doi:10.1038/s42255-024-01100-0 (2024).

70 Furlan, A. et al. Neurotensin neurons in the extended amygdala control dietary choice and energy homeostasis. Nature neuroscience 25, 1470–1480, doi:10.1038/s41593-022-01178-3 (2022).

71 Bolger, A. M., Lohse, M. & Usadel, B. Trimmomatic: a flexible trimmer for Illumina sequence data. Bioinformatics 30, 2114–2120, doi:10.1093/bioinformatics/btu170 (2014).

72 Dobin, A. et al. STAR: ultrafast universal RNA-seq aligner. Bioinformatics 29, 15–21, doi:10.1093/bioinformatics/bts635 (2013).

73 Liao, Y., Smyth, G. K. & Shi, W. featureCounts: an efficient general purpose program for assigning sequence reads to genomic features. Bioinformatics 30, 923–930, doi:10.1093/bioinformatics/btt656 (2014).

74 Robinson, M. D., McCarthy, D. J. & Smyth, G. K. edgeR: a Bioconductor package for differential expression analysis of digital gene expression data. Bioinformatics 26, 139–140, doi:10.1093/bioinformatics/btp616 (2010).

75 Yu, G., Wang, L. G., Han, Y. & He, Q. Y. clusterProfiler: an R package for comparing biological themes among gene clusters. OMICS 16, 284–287, doi:10.1089/omi.2011.0118 (2012).

76 Trubetskoy, V. et al. Mapping genomic loci implicates genes and synaptic biology in schizophrenia. Nature 604, 502–508, doi:10.1038/s41586-022-04434-5 (2022).

77 Farsi, Z. et al. Brain-region-specific changes in neurons and glia and dysregulation of dopamine signaling in Grin2a mutant mice. Neuron 111, 3378–3396 e3379, doi:10.1016/j.neuron.2023.08.004 (2023).

78 Langmead, B. & Salzberg, S. L. Fast gapped-read alignment with Bowtie 2. Nat Methods 9, 357-359, doi:10.1038/nmeth.1923 (2012).

79 Zhang, Y. et al. Model-based analysis of ChIP-Seq (MACS). Genome Biol 9, R137, doi:10.1186/gb-2008-9-9-r137 (2008).

80 Lun, A. T. & Smyth, G. K. csaw: a Bioconductor package for differential binding analysis of ChIP-seq data using sliding windows. Nucleic Acids Res 44, e45, doi:10.1093/nar/gkv1191 (2016).

81 Ramirez, F. et al. deepTools2: a next generation web server for deep-sequencing data analysis. Nucleic Acids Res 44, W160–165, doi:10.1093/nar/gkw257 (2016).

82 Gu, Z., Eils, R., Schlesner, M. & Ishaque, N. EnrichedHeatmap: an R/Bioconductor package for comprehensive visualization of genomic signal associations. BMC Genomics 19, 234, doi:10.1186/s12864-018-4625-x (2018).

