## Supplementary figures and images for "Rescue schizophrenia-related phenotypes caused by *Setd1a* deficiency by histone demethylase inhibitors"

### Extended Data Figure 1

**Extended Data Fig. 1**

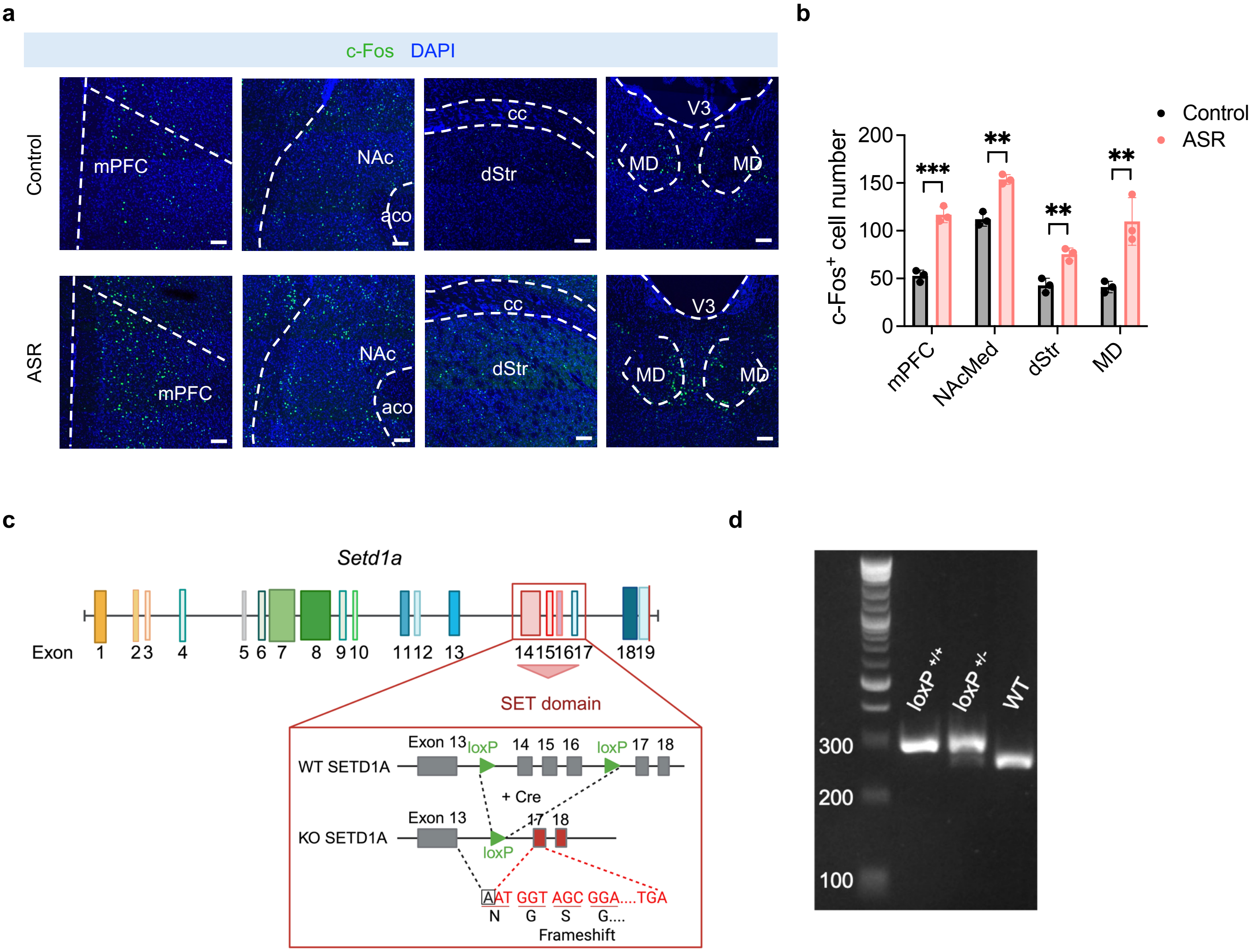

### Extended Data Figure 2

##### Extended Data Fig. 2

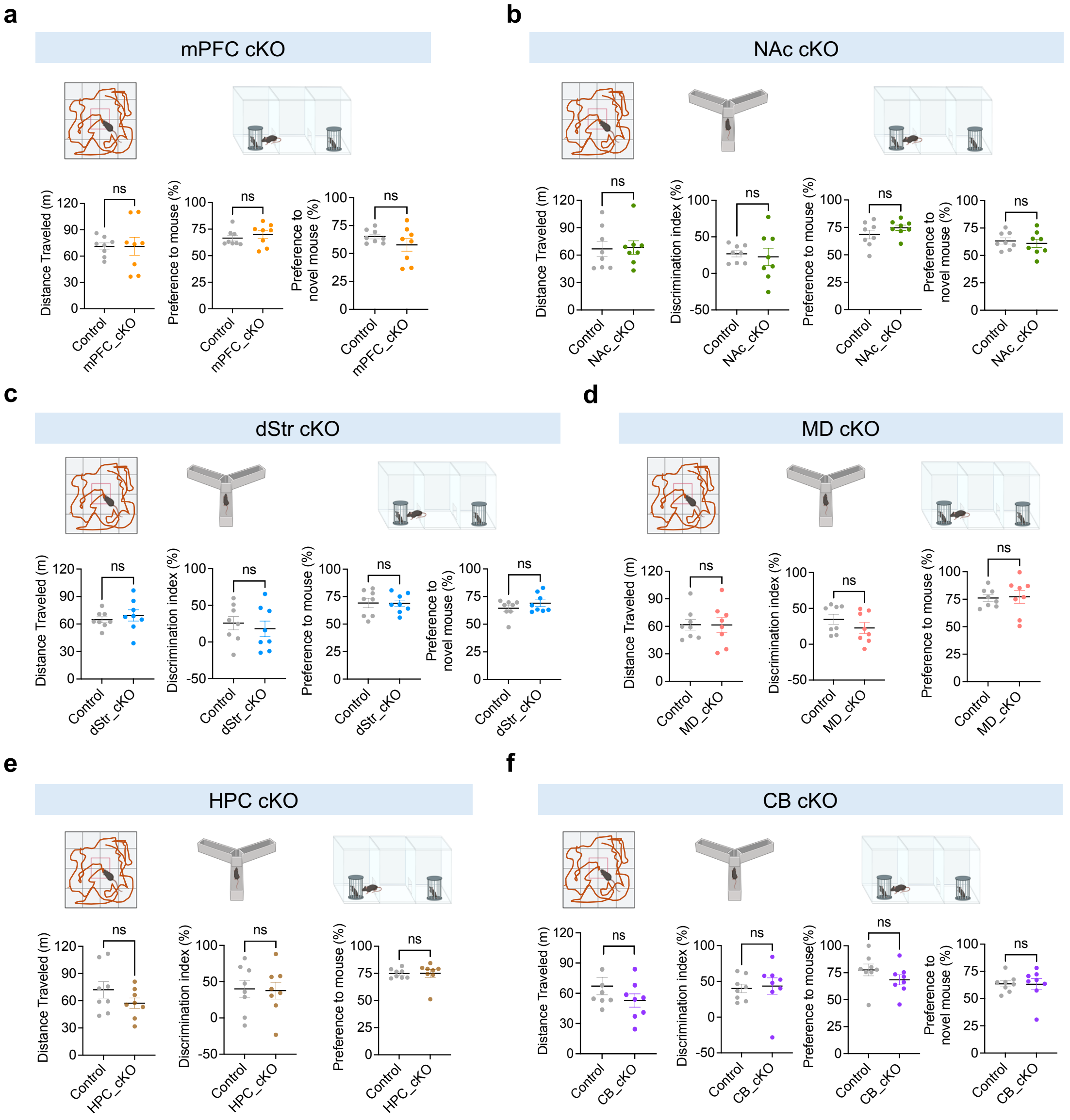

### Extended Data Figure 3

**Extended Data Fig. 3**

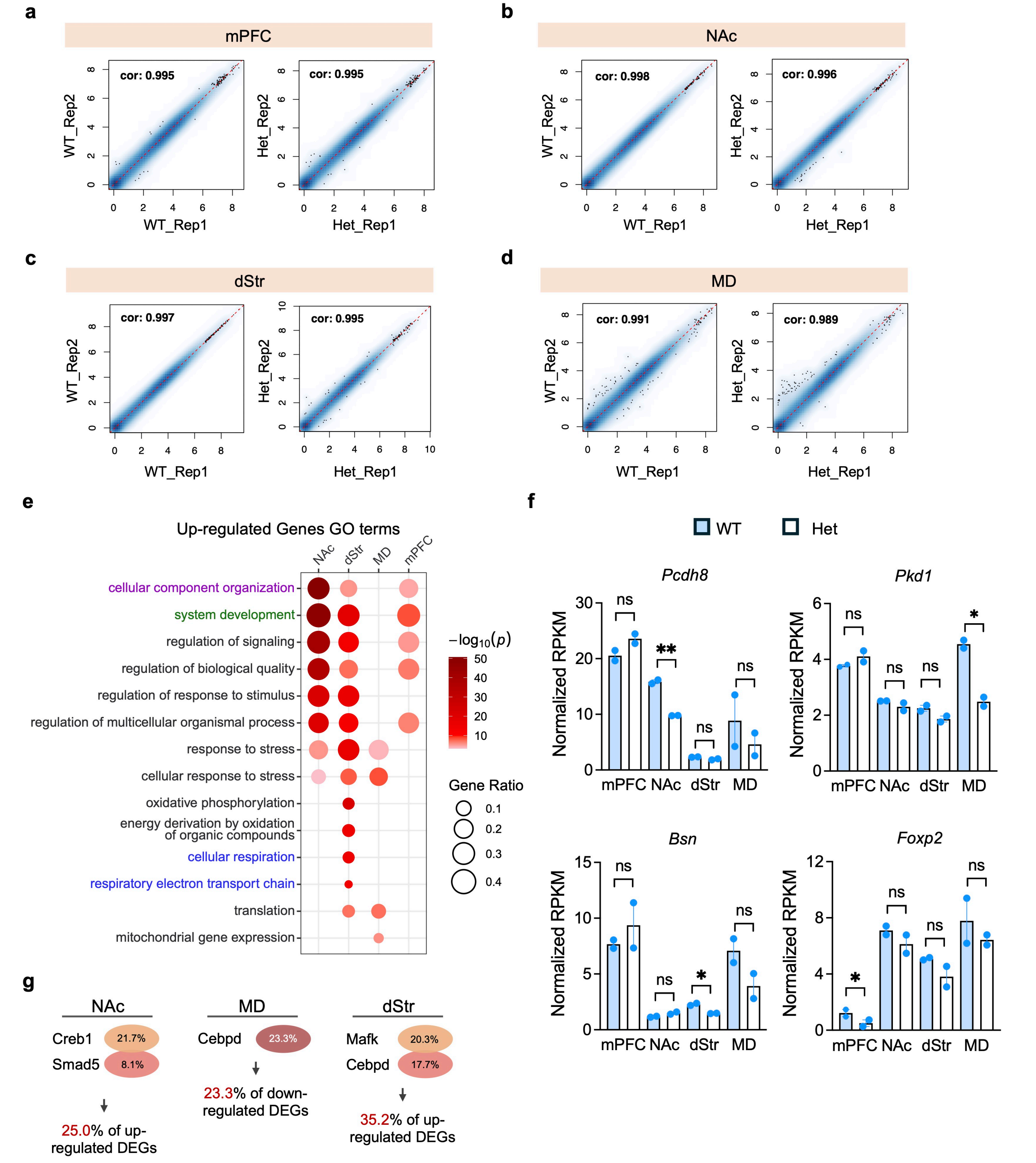

### Extended Data Figure 4

##### Extended Data Fig. 4

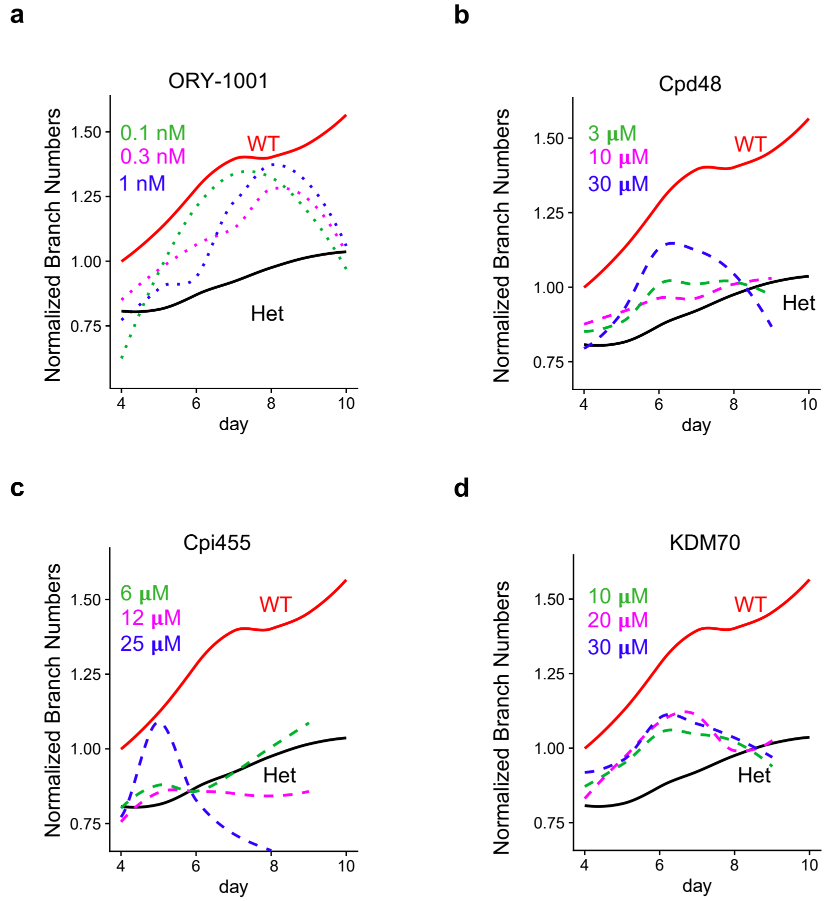

### Extended Data Figure 5

**Extended Data Fig. 5**

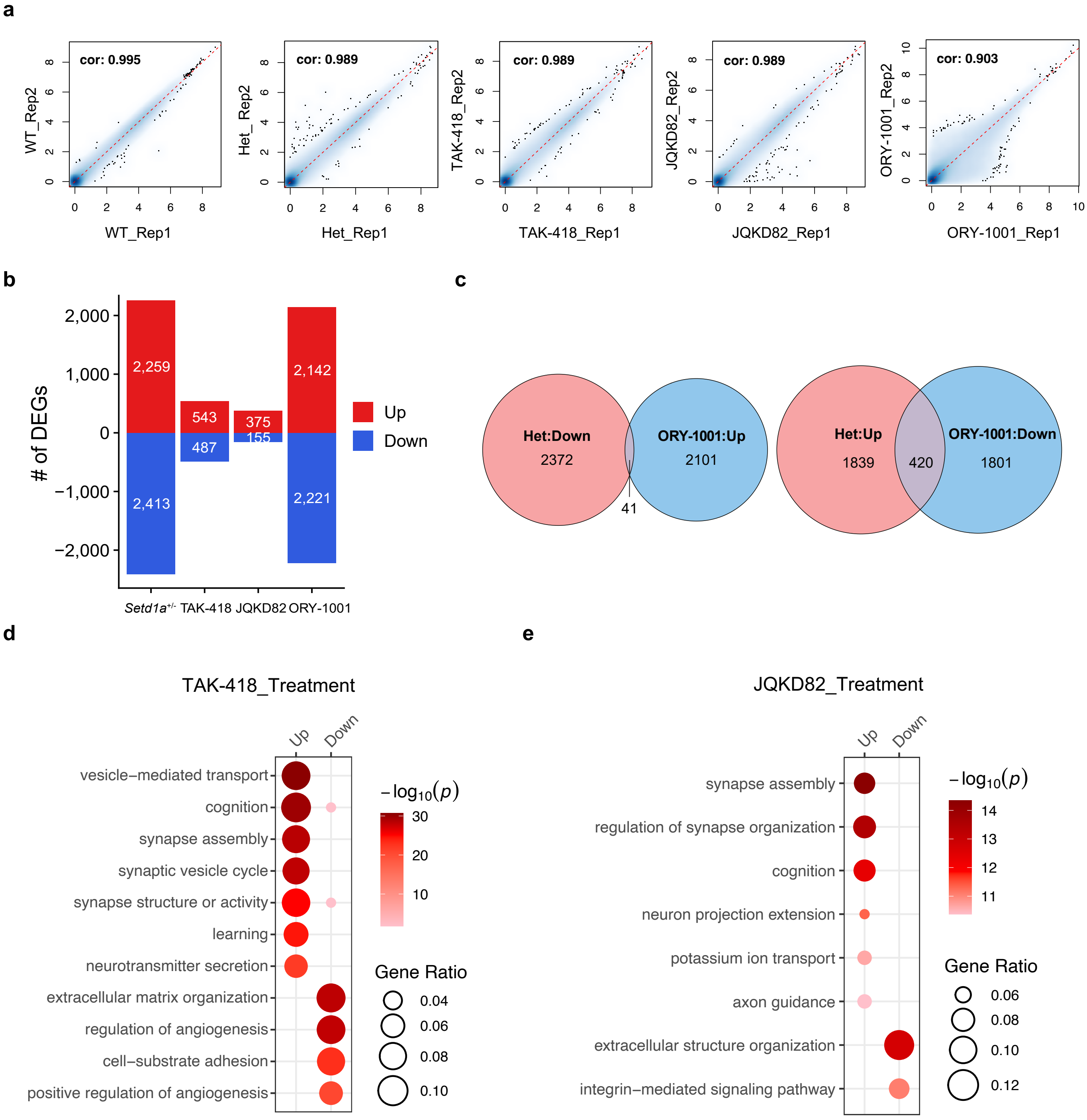

### Extended Data Figure 6

#### Extended Data Fig. 6

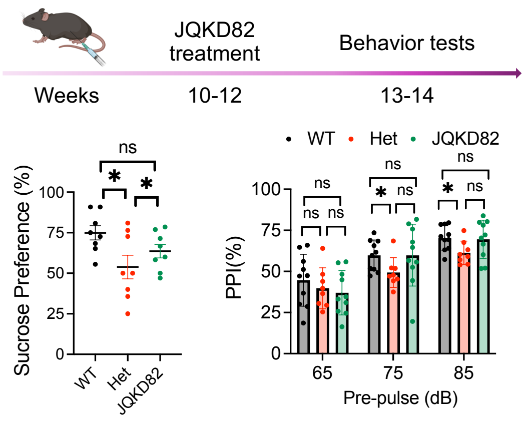

### Extended Data Figure 7

#### Extended Data Fig. 7

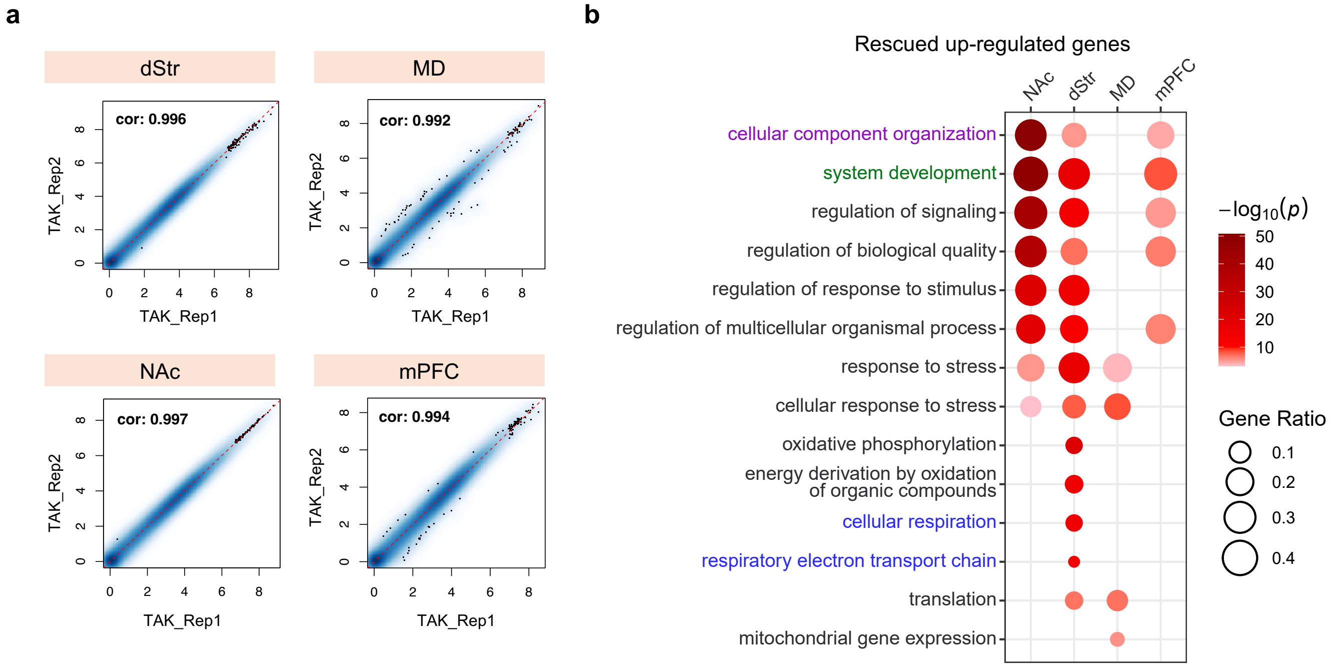

### Extended Data Figure 8

##### Extended Data Fig. 8

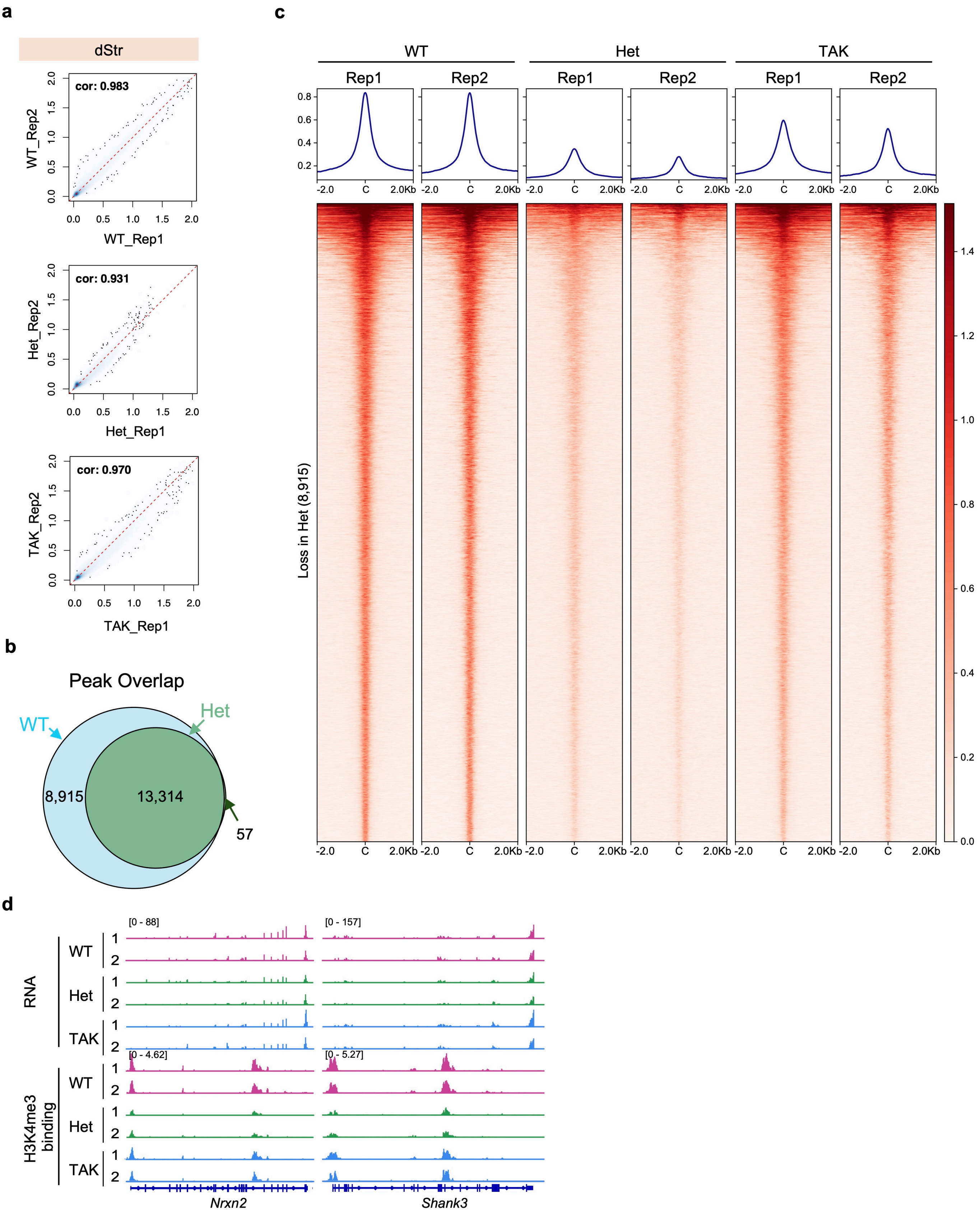
